# A conserved signaling pathway activates bacterial CBASS immune signaling in response to DNA damage

**DOI:** 10.1101/2022.04.27.489752

**Authors:** Rebecca K. Lau, Eray Enustun, Yajie Gu, Justin V. Nguyen, Kevin D. Corbett

## Abstract

To protect themselves from the constant threat of bacteriophage (phage) infection, bacteria have evolved diverse immune systems including restriction/modification, CRISPR/Cas, and many others. Here we describe the discovery of a two-protein transcriptional regulator module associated with hundreds of CBASS (<u>C</u>yclic oligonucleotide <u>B</u>ased <u>A</u>nti-phage <u>S</u>ignaling <u>S</u>ystem) immune systems, and demonstrate that this module drives expression of its associated CBASS system in response to DNA damage. We show that the helix-turn-helix transcriptional repressor CapH binds the promoter region of its associated CBASS system to repress transcription until it is cleaved by the metallopeptidase CapP. CapP is inactive except in the presence of single-stranded DNA, and CapP activity in cells is stimulated by DNA-damaging drugs. Together, CapH and CapP drive increased expression of their associated CBASS system in response to DNA damage. In both their structures and mechanisms, CapH and CapP resemble regulators that drive increased expression of DNA damage response genes in radiation-resistant *Deinococcus*, and control the mobilization of prophages and mobile elements in response to DNA damage. We also identify CapH and CapP-related proteins associated with diverse known and putative bacterial immune systems, including DISARM and two uncharacterized operons encoding proteins related to eukaryotic ubiquitin signaling pathways. Overall, our data highlight a mechanism by which bacterial immune systems can sense and respond to a universal stress signal, potentially enabling multiple immune systems to mount a coordinated defensive effort against an invading pathogen.

## Introduction

In all organisms, survival depends on the ability of cells to sense and respond to both internal and external threats. In addition to environmental stress, bacteria are continually challenged by bacteriophages (phages), and have evolved a wide array of immune systems to protect themselves from phage infection and propagation. Many anti-phage immune systems, including restriction-modification and CRISPR/Cas systems, specifically recognize and destroy foreign DNA to prevent phage replication (Mohanraju *et al*, 2016; Makarova *et al*, 2013). Other immune systems, termed abortive infection systems, sense phage infection and respond by killing the host cell, thereby preventing phage propagation and further infection in the bacterial community (Dy *et al*, 2014; Makarova *et al*, 2011a; Doron *et al*, 2018; Hampton *et al*, 2020). In many bacteria, multiple immune systems co-exist in so-called “defense islands” (Doron *et al*, 2018; Makarova *et al*, 2011b) and may cooperate, with non-lethal systems acting as a first line of defense and abortive infection systems becoming activated only as a last resort (Bernheim & Sorek, 2020).

The widespread and functionally-diverse CBASS anti-phage immune systems use an abortive infection mechanism in which a cGAS/DncV-like nucleotidyltransferase (CD-NTase) is activated upon phage infection and synthesizes a cyclic oligonucleotide secondary messenger (Ye *et al*, 2020; Lau *et al*, 2020; Cohen *et al*, 2019; Whiteley *et al*, 2019; Davies *et al*, 2012). This molecule in turn activates one of a variety of effector proteins, including phospholipases, nucleases, or pore-forming proteins to kill the host cell (Severin *et al*, 2018; Lowey *et al*, 2020; Lau *et al*, 2020; Cohen *et al*, 2019). While so-called Type I CBASS systems encode only a CD-NTase and a cell-killing effector protein, the majority of CBASS systems encode ancillary genes putatively involved in infection sensing and/or CD-NTase activation (Millman *et al*, 2020; Burroughs *et al*, 2015). Type II CBASS systems encode two proteins, Cap2 and Cap3, that are related to eukaryotic ubiquitination machinery and are required for protection against phage (Cohen *et al*, 2019; Ledvina *et al*, 2022). Type III CBASS systems, meanwhile, encode peptide-binding HORMA domain proteins (Cap7 and Cap8) that are proposed to bind specific peptides to sense infection and then activate their associated CD-NTase (Ye *et al*, 2020).

All CBASS systems are thought to directly sense phage infection and respond by triggering cell death. Here, we identify a pair of transcriptional regulators, termed CapH and CapP, that are associated with hundreds of CBASS systems and upregulate CBASS expression in response to DNA damage. DNA damage is a universal stress signal in bacterial cells (Benler & Koonin, 2020), and we show that CapH and CapP are structurally and functionally similar to regulators that mobilize prophages and integrative and conjugative elements (ICE elements), and to activators of the DNA damage response in radiation-resistant *Deinococcus* species. We also identify CapH and CapP-like regulators associated with a variety of known or putative bacterial immune systems, revealing that these proteins represent a conserved signaling module that regulates defense-system expression in response to DNA damage across bacteria.

## Results

### Identification of *capH* and *capP* genes associated with CBASS systems

We previously showed that a Type III CBASS system from *E. coli* strain MS115-1 provides robust protection against bacteriophage λ through an abortive infection mechanism (Ye *et al*, 2020; Lau *et al*, 2020). Examination of this system’s genomic neighborhood revealed a pair of genes directly upstream of the core CBASS genes and encoded on the opposite strand (i.e. sharing a promoter region with the core CBASS genes), that encode a predicted helix-turn-helix (HTH) DNA binding protein and a predicted Zn^2+^ metallopeptidase (**Figure 1A**). We term these two genes *capH* (<u>C</u>BASS <u>a</u>ssociated <u>p</u>rotein, <u>H</u>elix-turn-helix) and *capP* (<u>C</u>BASS <u>a</u>ssociated <u>p</u>rotein, <u>P</u>eptidase). In their position and orientation relative to the core CBASS genes, *capH* and *capP* are similar to *capW*, a transcriptional regulator associated with a distinct subset of CBASS systems (Blankenchip *et al*, 2022). BLAST searches revealed that CapP shares strong similarity to IrrE, a metallopeptidase that regulates the DNA damage response in *Deinococcus* by cleaving an HTH-family transcription factor, DdrO (Ludanyi *et al*, 2014; Vujičić-Žagar *et al*, 2009). DdrO normally binds the promoters of DNA damage response genes and suppresses their expression, but upon DNA damage IrrE becomes activated and cleaves DdrO, releasing it from DNA and activating expression of the DNA damage response genes (Blanchard *et al*, 2017; Ludanyi *et al*, 2014; de Groot *et al*, 2019). The similarity of CapP to IrrE, and its association with the HTH protein CapH, suggested that CapH and CapP may functionally cooperate to control expression of their associated CBASS system.

**Figure 1.**
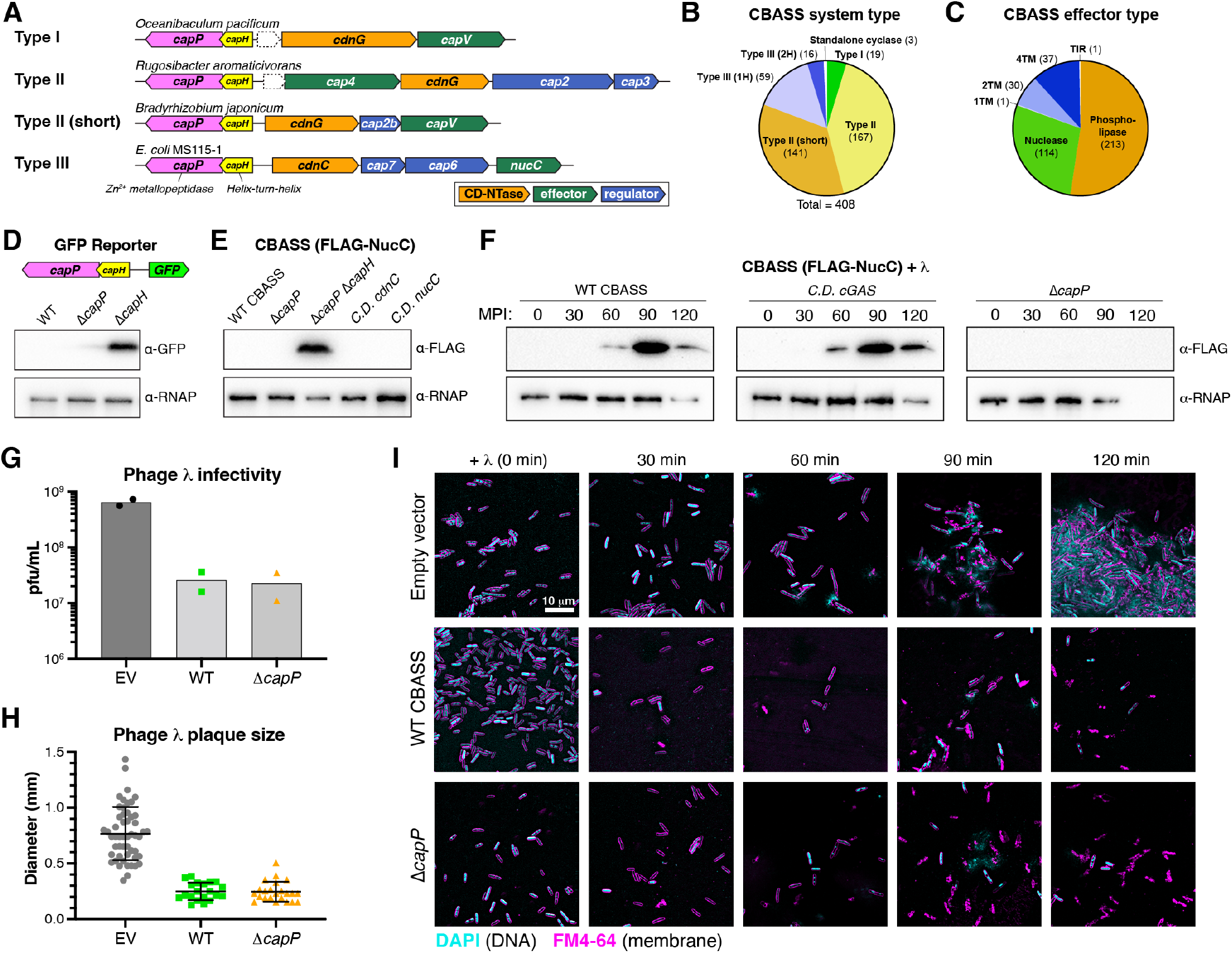
Identification of CBASS-associated genes *capH* and *capP* and role of CapH and CapP in the CBASS antiviral response. (A) Operon schematics of four representative CBASS systems with associated *capH* (yellow) and *capP* (pink) genes. See **Table S1** for all identified systems. For each system, core CBASS genes are colored as in the key: CD-NTases orange, putative regulator(s) blue, and effector(s) green. Dotted outlines indicate unknown theoretical genes. (B) Distribution of *capH*+*capP*-associated CBASS systems, sorted by system type as defined by Millman et al. (Millman *et al*, 2020). Type III (1H) and Type III (2H) refer to Type III systems with one or two HORMA domain proteins, respectively. (C) Distribution of *capH*+*capP*-associated CBASS systems, sorted by effector type as defined by Millman et al. (Millman *et al*, 2020). 1TM, 2TM, and 4TM refer to effectors with one, two, or four predicted transmembrane segments, respectively. (D) *Top:* Schematic of GFP expression reporter system, with the CBASS promoter, *capH*, and *capP* genes from *E. coli* MS115-1 and the CBASS core genes replaced with GFP. *Bottom:* Western blot showing GFP expression in cells with the wild-type GFP reporter or constructs lacking either *capP* or *capH* genes. RNAP: RNA polymerase loading control. See full blot in **Figure S1A**. (E) Western blot showing FLAG-NucC expression in uninfected cells of the indicated genotype. C.D. *cdnC*: D72N/D74N catalytic dead mutant; C.D. *nucC*: D73N catalytic dead mutant. RNAP: RNA polymerase loading control. See full blot in **Figure S1B**. (F) Western blots of the CBASS expression reporter system with FLAG-NucC, showing FLAG-NucC expression after infection with phage λ cI- (multiplicity of infection: 10). MPI: minutes post infection. α-RNAP: anti-RNA polymerase loading control. Low RNAP expression at later time points is due to cell death. (G) Quantitative plaque assay showing infectivity of λ cI- against cells containing no CBASS system (EV: empty vector), the wild-type *E. coli* MS115-1 CBASS systems (WT), or a mutant system lacking *capP* (Δ*capP*). Data is shown in units of plaque forming units per mL of purified phage (pfu/mL). (H) Size of phage plaques for λ cI-infecting cells containing no CBASS system (EV: empty vector), the wild-type *E. coli* MS115-1 CBASS systems (WT), or a mutant system lacking *capP* (Δ*capP*). (I) Live-cell fluorescence microscopy of λ cI-infecting cells containing no CBASS system (EV: empty vector), the wild-type *E. coli* MS115-1 CBASS systems (WT), or a mutant system lacking *capP* (Δ*capP*).

We systematically searched the genomic neighborhoods of 6,233 bacterial CBASS CD-NTases (Cohen *et al*, 2019) for genes related to *capP* and identified 408 CBASS systems with a predicted Zn^2+^ metallopeptidase within 10 kb of the system’s CD-NTase gene. We manually inspected each system and found that the vast majority encode both *capH* and *capP* genes upstream of the core CBASS genes and on the opposite strand (**Figure 1A, Table S1**). In these systems, *capP* is most often annotated as a “domain of unknown function” (DUF) 955 or PFAM06114 protein. We identified *capH* and *capP* genes associated with Type I, Type II, and Type III CBASS systems that encode a variety of predicted effectors including phospholipases, transmembrane proteins, and endonucleases (**Figure 1B-C, Table S1**). In 24 systems, *capH* and *capP* are encoded alongside a predicted σ70-family σ factor (**Table S1**), while 15 systems encode an apparent fusion of *capH* and *capP* (**Table S1**).

To determine whether *capH* and *capP* control expression of their associated CBASS operon, we generated a reporter construct with *capH*, *capP*, and the promoter region of the CBASS system from *E. coli* MS115-1, plus a gene encoding GFP (green fluorescent protein) in place of the core CBASS genes. When both *capH* and *capP* were present, expression of GFP in uninfected log-phase cells was too low for detection by anti-GFP immunoblotting (**Figure 1D**). Expression was also nearly undetectable in a strain lacking *capP*, but we observed high GFP expression in a strain lacking *capH* (**Figure 1D, S1A**). We confirmed these findings using a separate reporter construct encoding the full six-gene CBASS system of *E. coli* MS115-1, with a N-terminal FLAG tag fused to the effector nuclease *nucC* (**Figure 1E, S1B**). Anti-FLAG immunoblots showed that basal *nucC* expression is undetectable in the wild-type CBASS system, but that expression is dramatically boosted upon deletion of both *capH* and *capP* (**Figure 1E**). Overall, these data suggest that in uninfected cells, CapH acts as transcriptional repressor for its associated CBASS system, similar to the repression of DNA damage response genes by *Deinococcus* DdrO.

### CapP controls CBASS expression but is not required for phage protection

Our reporter assays indicated that CapH acts as a transcriptional repressor for its associated CBASS operon. To determine the role of CapP in CBASS expression, we first used our FLAG-*nucC* reporter system to test for CBASS expression changes upon infection with an obligately lytic variant of bacteriophage λ lacking the cI gene (λ cI^-^) (Rajagopala *et al*, 2011). With the wild-type CBASS system encoding *capH* and *capP*, we observed a strong increase in FLAG-NucC expression starting ∼60 minutes after infection and peaking around 90 minutes after infection (**Figure 1F**). In the absence of *capP*, we observed no such increase in FLAG-NucC expression (**Figure 1F**). We also observed increased FLAG-NucC expression in a system with catalytically-dead CD-NTase (CdnC D72N/D74N), indicating that the observed expression changes do not depend on CBASS signaling (**Figure 1F**). These data suggest that CapP responds to phage infection by antagonizing CapH, resulting in a loss of repression and an increase in CBASS expression.

To test the role of CapH and CapP in phage protection, we compared the ability of wild-type *E. coli* MS115-1 CBASS and a mutant lacking *capP* to protect against λ cI^-^ infection. We previously reported that when cloned into an IPTG-inducible expression vector, the four core genes from *E. coli* MS115-1 CBASS (*cdnC* (CD-NTase), *cap7* (HORMA), *cap6* (TRIP13), and *nucC*) provide strong protection against λ cI^-^ (Ye *et al*, 2020; Lau *et al*, 2020). We found that the native six-gene operon also encoding *capH* and *capP* also provides protection against λ cI^-^, as measured by both a reduction in viral plaque numbers (**Figure 1G**) and a reduction in plaque size (**Figure 1H**) compared to bacteria lacking CBASS. Notably, a system lacking *capP* provided protection equivalent to the wild-type system (**Figure 1G-H**), indicating that the boost in CBASS expression we observe upon λ cI^-^ infection is not required for the system’s antiviral function. This conclusion was supported by live-cell microscopy: cells lacking CBASS were lysed and showed significant extracellular DNA 90-120 minutes after infection, while cells with either wild-type CBASS or a system lacking *capP* showed strong degradation of cellular DNA within 30-60 minutes of infection (**Figure 1I**). These data are consistent with our previous finding that activation of the NucC nuclease results in DNA degradation and host-cell death (Lau *et al*, 2020). Thus, *capP* is not required for CBASS activation or protection against the lytic phage λ cI^-^ in the *E. coli* MS115-1 CBASS system, leaving open the question of its role in CBASS signaling.

### CapH binds the promoter region of its associated CBASS system

Our reporter assays suggested that CapH acts as a transcriptional repressor for its associated CBASS system, potentially by directly binding the CBASS promoter region. To further explore this idea, we first cloned and purified the protein’s predicted helix-turn-helix (HTH) DNA binding domain (residues 2-67; **Figure 2A**), and determined a 1.02 Å-resolution crystal structure of this domain (**Figure 2B, Table S2**). The domain forms a canonical HTH fold, and modeling a DNA-bound complex based on known HTH-DNA complexes revealed several highly-conserved residues putatively involved in DNA binding, including Ser32, Arg40, and Arg44 (**Figure 2B**). To identify the binding site for CapH, we searched the promoter region of its associated CBASS system for conserved elements near the transcription start site. We identified a highly conserved 22-bp region upstream of the promoter’s predicted -35 site (**Figure 2C**), and found that CapH robustly binds this sequence (*K_d_* = 0.6 µM) while showing no binding to a random control sequence (**Figure 2D**). We found that mutation of Ser32 or Arg44 on the predicted DNA-binding face of CapH eliminated detectable DNA binding, while mutation of Arg40 reduced, but did not eliminate, DNA binding (**Figure 2E**). We next tested the same mutations in our GFP reporter system, and found that mutation of Ser32 resulted in high expression of GFP (**Figure 2F**). These data are consistent with CapH acting as a transcriptional repressor for its associated CBASS operon.

**Figure 2.**
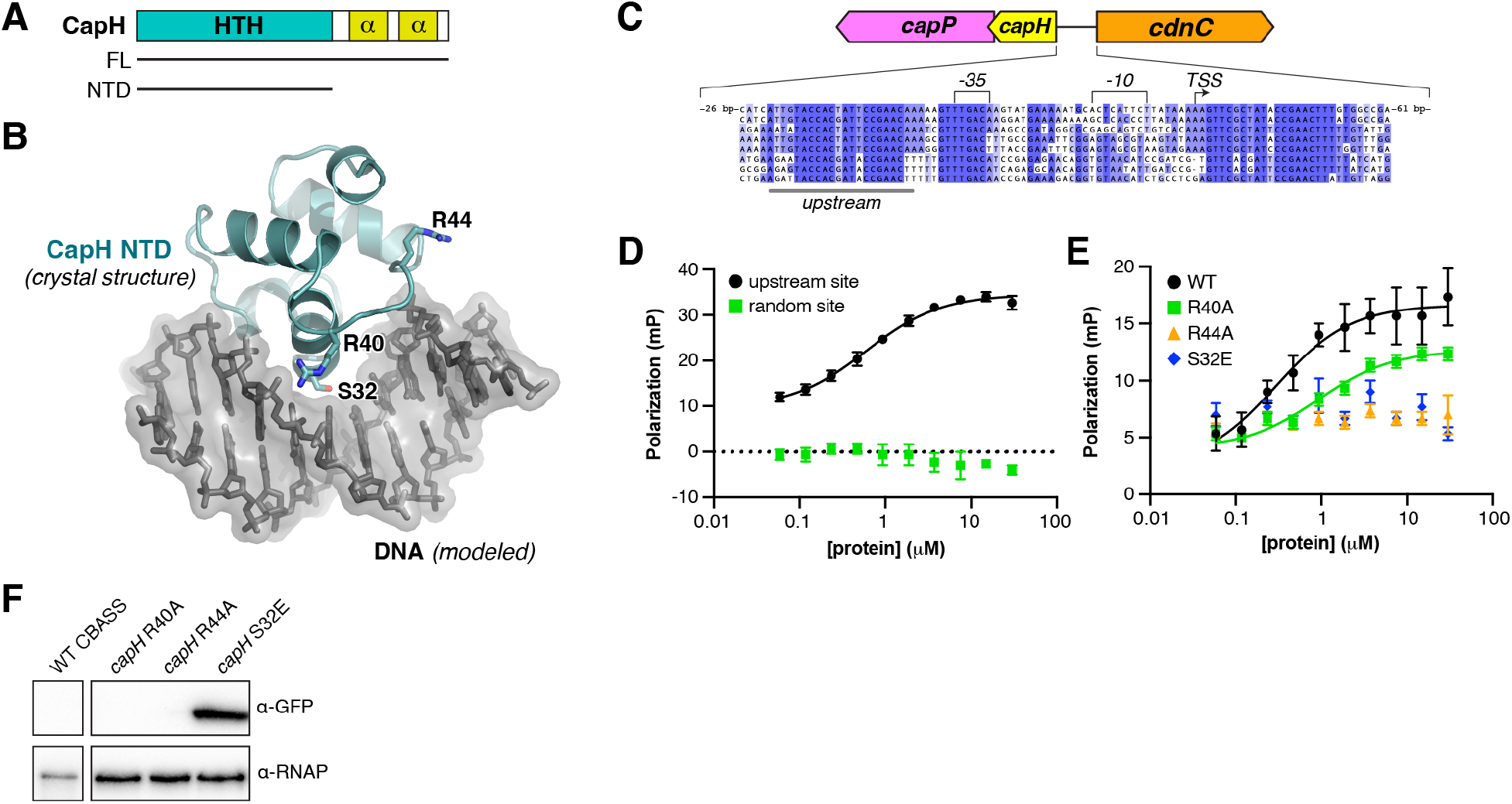
CapH binds the CBASS promoter region. (A) Domain schematic of *E. coli* MS115-1 CapH, and truncation used for crystallization of the N-terminal HTH domain comprised of residues 2-67 (NTD). (B) Crystal structure of the CapH NTD (blue), with bound DNA modeled from a structural overlay with a known HTH-DNA complex structure (PDB ID 3CLC; (McGeehan *et al*, 2008)). Shown in sticks are three conserved residues (S32, R40, and R44) putatively involved in DNA binding. (C) Sequence alignment of CBASS promoter regions in *E. coli* MS115-1 (NCBI RefSeq GG771785), *Cronobacter sakazakii* strain cro3915C2 (NZ_NRJY01000012), *Pseudomonas stutzeri* strain KC NODE_1_length_951488_cov_16.453 (NZ_POUN01000001), *Pseudomonas* sp. RIT 412 RIT412_S3_7 (NZ_QBJA02000007), *Pseudomonas* sp. MF4836 (NZ_MVOL01000002), *Burkholderia pseudomallei* strain MSHR4301 (NZ_LXCN01000015), *Ralstonia insidiosa* strain WCHRI065162 (NZ_PKPC01000011), *Thauera* sp. K11 plasmid pTX1 (NZ_CP023440). Promoter sequences (-35, -10, and TSS) were predicted by the BPROM server (Salamov & Solovyevand, 2011). The conserved upstream region is noted with a gray underline. (D) Fluorescence polarization assay showing binding of *E. coli* MS115-1 CapH to a 22-bp DNA containing the upstream site from panel (A) (black; *K_d_*=0.6 +/- 0.1 μM), and a random 22-bp DNA (green) (E) Fluorescence polarization assay showing binding of *E. coli* MS115-1 CapH (wild-type or indicated point mutants) to the 22-bp upstream site DNA. WT *K_d_*=0.3 +/- 0.1 μM; R40A *K_d_* =0.9 +/- 0.3 μM; no binding detected for R44A or S32E). (F) GFP expression reporter assay showing loss of suppression upon mutation of CapH. RNAP: RNA polymerase loading control. See full blot in **Figure S1A**.

### CapH oligomerization is required for DNA binding

In the *Deinococcus* DdrO-IrrE system, DdrO forms a homodimer through its C-terminal domain and this dimerization is required for DNA binding and transcriptional repression by the protein (de Groot *et al*, 2019; Ludanyi *et al*, 2014). To test whether CapH forms an oligomer, we used size exclusion chromatography coupled to multi-angle light scattering (SEC-MALS). We found that full-length CapH forms an oligomer with an overall size consistent with either a homotrimer or a mixture of dimer and tetramer states (**Figure 3A-B**). The isolated C-terminal region of CapH (residues 67-107; CapH^CTD^) forms a similar oligomer, while the isolated N-terminal HTH domain is monomeric (**Figure 3B**). These data show that CapH oligomerizes through its C-terminal domain. To determine the mechanism of oligomerization, we crystallized and determined a 1.75 Å-resolution crystal structure of CapH^CTD^ (**Figure 3C, Table S2**). In the structure, four CapH^CTD^ protomers form a tight homotetramer with a dimer-of-dimers architecture. Each CapH^CTD^ protomer forms two α-helices that fold into a V shape, with the homodimer assembled by two protomers arranged antiparallel to one another with the V shapes interlocked. The CapH^CTD^ homodimer is stabilized by a hydrophobic core comprising Phe81, Tyr85, Leu96, and Leu100 of each protomer (**Figure 3C**). The CapH^CTD^ homodimer resembles the C-terminal dimerization domains of other dimeric bacterial transcription factors, including *Mycobacterium tuberculosis* EspR, *Bacillus subtilis* CinR, and *Citrobacter* C.Csp231I (Shevtsov *et al*, 2015; Lewis *et al*, 1998; Gangwar *et al*, 2014). The CapH^CTD^ homotetramer is assembled through a separate hydrophobic interface between the C-terminal α-helices of four protomers, involving residues Ile99 and Phe103 (**Figure 3C**).

**Figure 3.**
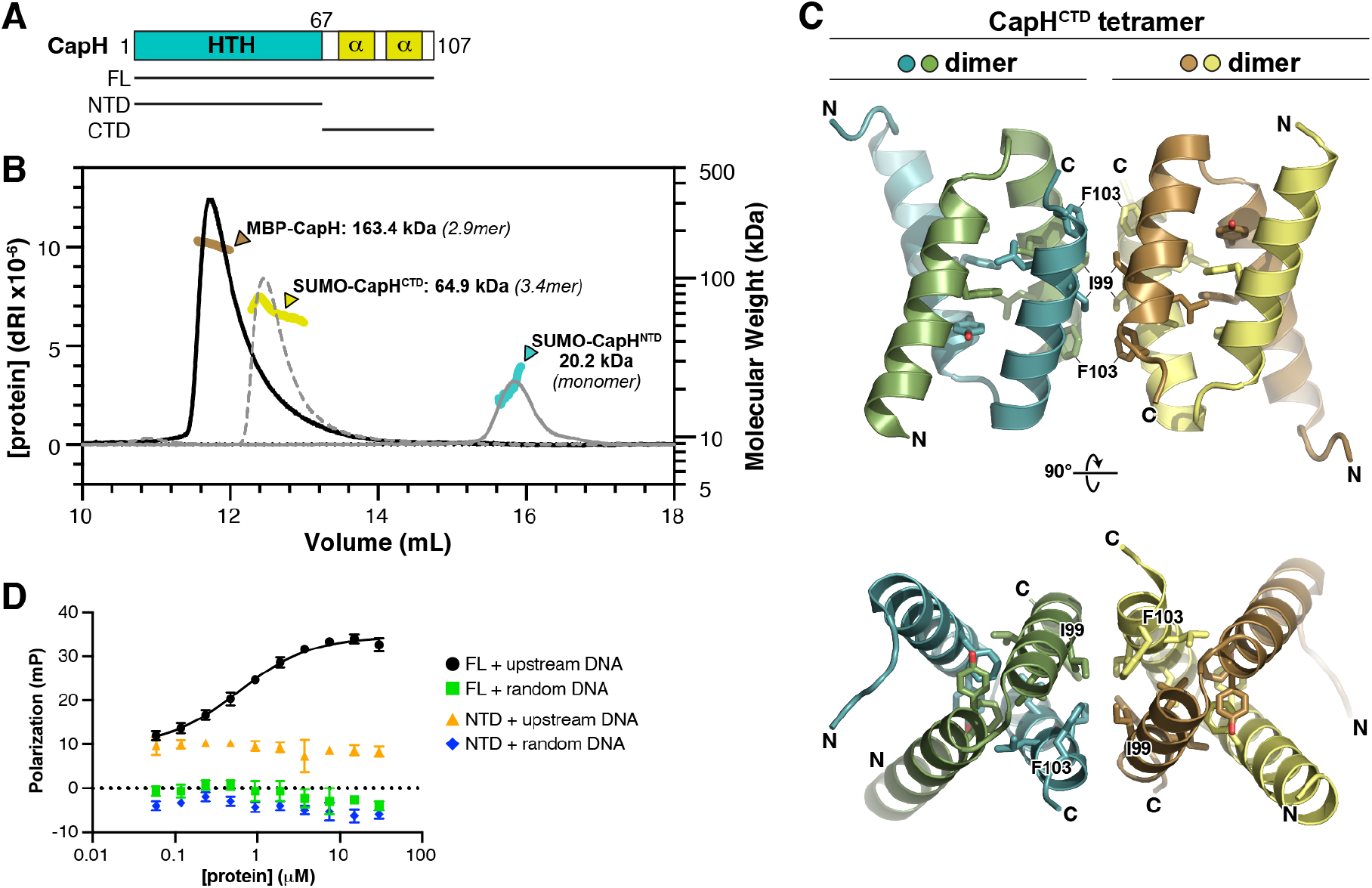
CapH oligomerization is required for DNA binding. (A) Domain schematic of *E. coli* MS115-1 CapH, showing the truncations used for oligomeric state determination. (B) Size exclusion chromatography coupled to multi-angle light scattering (SEC-MALS) determination of CapH oligomeric state. For each construct, measured molecular weight and inferred oligomer are indicated. MBP-fused full-length CapH (monomer MW=57.0 kDa) is shown in brown; SUMO-fused CapH^CTD^ (monomer MW=19.1 kDa) is yellow, and SUMO-fused CapH^NTD^ (monomer MW=22.0 kDa) is cyan. (C) Crystal structure of the CapH^CTD^ homotetramer. Residues comprising the hydrophobic core of each dimer (Phe81, Tyr85, Leu96, and Leu100) are shown as sticks, and residues comprising the hydrophobic tetramerization interface (Ile99 and Phe103) are shown as sticks and labeled. See **Figure S2A** for structure of the CapH^CTD^(I99M) homodimer. (D) Fluorescence polarization assay showing binding of *E. coli* MS115-1 CapH (full-length or NTD) to the 22-bp upstream site or random DNA. WT *K_d_*=0.6 +/- 0.1 μM; no binding detected for CapH^NTD^). See **Figure S2C** for DNA binding of CapH(I99M).

During the course of structure determination for CapH^CTD^, we generated a construct with a mutation of Ile99 to methionine (CapH^CTD^(I99M)). We determined a 1.26 Å-resolution structure of this mutant, revealing a CapH homodimer equivalent to our structure of wild-type CapH^CTD^, but lacking the tetrameric assembly (**Figure S2A**). Consistent with this finding, SEC-MALS showed that CapH(I99M) forms a stable homodimer in solution, rather than the larger oligomer observed with wild-type CapH (**Figure S2B**). Thus, the I99M mutant disrupts CapH tetramerization, but not dimerization.

We next tested the role of CapH oligomerization in DNA binding. We used fluorescence polarization to compare the DNA binding affinity of full length wild-type CapH to that of the CapH HTH domain (residues 2-67; CapH^NTD^), which forms a monomer; and to the CapH(I99M) mutant, which forms a homodimer. We observed no binding of CapH^NTD^ to DNA, demonstrating that CapH oligomerization is required for DNA binding (**Figure 3D**). The affinity of CapH(I99M) for DNA (*K_d_* = 2.3 μM) is about 10-fold lower than wild-type CapH (*K_d_* = 0.3 μM), demonstrating that tetramerization of CapH contributes to high-affinity DNA binding (**Figure S2C**). In our GFP reporter system, however, CapH(I99M) effectively suppressed expression of GFP (**Figure S1A**), demonstrating that tetramer formation is not required for CapH function in cells. Thus, while dimerization of CapH is critical for its DNA binding activity, tetramerization is dispensable for both DNA binding and effective suppression of CBASS gene expression.

### The structure of CapP reveals an internal cysteine switch

In the *Deinococcus* DNA damage response pathway, cleavage of DdrO by the metallopeptidase IrrE results in loss of DNA binding by DdrO, enabling increased expression of DNA damage response genes (Ludanyi *et al*, 2014; de Groot *et al*, 2019). Our data on DNA binding and CBASS repression by CapH, and in particular the importance of CapH oligomerization for DNA binding, suggests a functional parallel between DdrO-IrrE and CapH-CapP. To better understand this relationship, we purified and determined a 1.35 Å-resolution crystal structure of CapP from a CBASS system in *Thauera* sp. K11 (56% identical to *E. coli* MS115-1 CapP; **Figure S3A**). The overall structure of CapP resembles that of IrrE, with the protein folding into three domains: an N-terminal Zn^2+^ metallopeptidase domain, a central linker domain with topology resembling a helix-turn-helix domain, and a C-terminal GAF domain (**Figure 4A-B**). The N-terminal domain closely resembles other HExxH Zn^2+^ metallopeptidases including IrrE, with five α-helices and a three-stranded β-sheet. A Zn^2+^ ion is coordinated in the conserved active site by residues His96, His100, and Glu129 (**Figure 4C-E**). The predicted active-site glutamate residue, Glu97, is positioned close by but not directly coordinating the bound Zn^2+^ ion. Instead, a conserved cysteine residue, Cys113, completes the coordination of the bound Zn^2+^ ion. Cys113 is located on an insertion in the metallopeptidase domain, on a β-strand that drapes over the active site in the same position that substrate peptides bind in related Zn^2+^ metallopeptidases (Cerdà-Costa & Xavier Gomis-Rüth, 2014). We term this region the cysteine switch loop, after the cysteine switch motif found in matrix metalloproteases. These enzymes are synthesized as inactive precursors with an N-terminal domain bearing a conserved cysteine residue (the cysteine switch) that coordinates the active-site Zn^2+^ ion and inhibits activity (**Figure S3B-C**). The protease is only activated upon proteolytic cleavage and dissociation of the cysteine switch domain (Van Wart & Birkedal-Hansen, 1990; Cerdà-Costa & Xavier Gomis-Rüth, 2014; Springman *et al*, 1990). Of the 408 CapP proteins associated with CBASS systems, 134 (33%) possess the cysteine switch loop, including *E. coli* MS115-1 CapP (**Figure 4F, S3D**).

**Figure 4.**
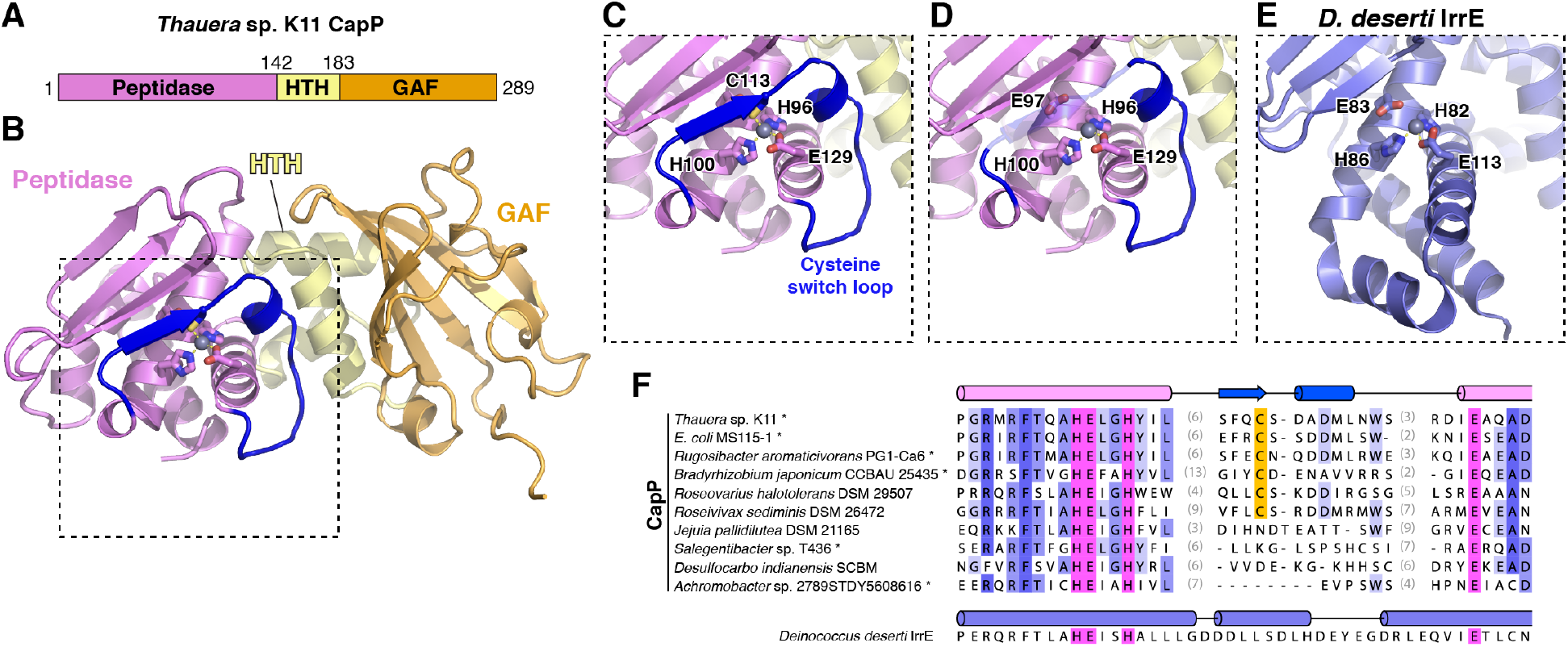
Structure of CapP reveals an internal cysteine switch. (A) Domain schematic of *Thauera* sp. K11 CapP, with N-terminal Zn^2+^ peptidase domain pink, central HTH domain yellow, and C-terminal GAF domain orange. See **Figure S3A** for comparison of the *Thauera* sp. K11 and *E. coli* MS115-1 CBASS operons. (B) Structure of *Thauera* sp. K11 CapP with domains colored as in panel (A). Shown in sticks are active-site residues H96, E97, H100, and E129, and a bound Zn^2+^ ion is shown as a gray sphere. Shown in blue is the internal cysteine switch loop, with C113 (shown as sticks) coordinating the bound Zn^2+^ ion. (C) Closeup view of the *Thauera* sp. K11 CapP active site. See **Figure S3B-C** for comparison with a cysteine switch-containing matrix metalloprotease. (D) Closeup view of the *Thauera* sp. K11 CapP active site as in panel (C), with transparent cysteine switch loop to show the position of the catalytic glutamate residue E97. (E) Equivalent closeup view of the *Deinococcus deserti* IrrE active site (PDB ID 3DTI; (Vujičić-Žagar *et al*, 2009)), showing the active site residues and bound Zn^2+^ ion. (F) Sequence alignment of representative CBASS-associated CapP proteins, showing the cysteine switch loop present in a subset of these proteins. See **Table S1** for full list, and **Figure S3D** for evolutionary tree of CapP with presence/absence of the cysteine switch annotated.

The presence of the cysteine switch loop in CapP suggests that the protein’s peptidase activity is tightly controlled, perhaps by a conformational change that induces dissociation of the cysteine switch from the CapP active site and allows substrate binding. In *Deinococcus* IrrE, the protein’s peptidase activity is thought to be activated upon DNA damage by the binding of an unknown ligand to the protein’s C-terminal GAF domain, although IrrE does not contain a cysteine switch loop (Vujičić-Žagar *et al*, 2009). In other proteins, GAF domains are known to bind nucleotide-based second messengers including cyclic GMP, which binds the GAF domain of phosphodiesterase 6C and allosterically regulates its enzymatic activity (Ho *et al*, 2000; Gross-Langenhoff *et al*, 2006; Levdikov *et al*, 2009; Martinez *et al*, 2002, 2008). When we compared the structure of CapP to that of cyclic GMP-bound phosphodiesterase 6C, we observed that CapP possesses a large number of surface-exposed aromatic residues near the putative ligand-binding site (**Figure S4A**). If CapP is allosterically regulated through the GAF domain, these residues may be involved in the binding of nucleotide-based ligand(s).

### CapP cleaves CapH when stimulated by single-stranded DNA

We next sought to directly test whether CapP cleaves CapH. Our initial tests using purified proteins in vitro showed no CapP-mediated cleavage of CapH (not shown), so we instead developed an assay to detect CapP activity in *E. coli* cells. We co-expressed CapP with a fusion protein comprising CapH with an N-terminal maltose binding protein (MBP) tag and a C-terminal GFP tag, and used an anti-GFP western blot to detect CapP activity. We detected a minor band representing a CapP cleavage product in the presence of wild-type CapP, but not when the CapP active site was mutated (Glu98 to glutamine; E98Q) (**Figure 5A**). At ∼32 kDa, this band likely represents a product of CapP cleavage near the C-terminus of CapH.

**Figure 5.**
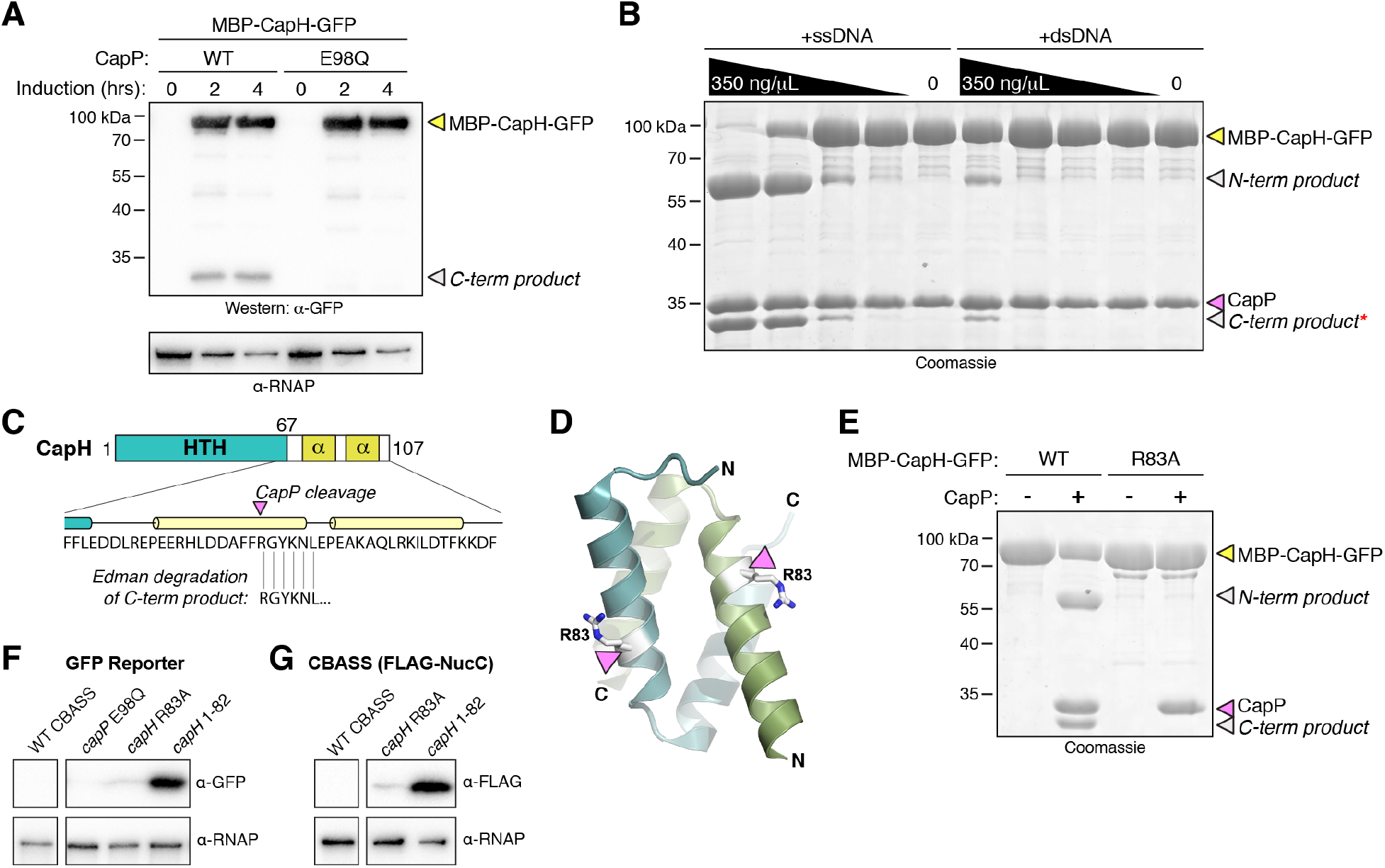
CapP cleaves CapH and is stimulated by single-stranded DNA. (A) Anti-GFP western blot showing coexpression in *E. coli* of an MBP-CapH-GFP fusion construct with wild-type or catalytic-dead (E98Q) CapP. Full-length MBP-CapH-GFP is indicated with a yellow arrowhead, and the C-terminal product of CapP cleavage is indicated with a white arrowhead. RNAP: RNA polymerase loading control. (B) In vitro cleavage of purified MBP-CapH-GFP (yellow arrowhead) into N-terminal and C-terminal products (white arrowheads) by CapP is stimulated by DNA. For both ssDNA and dsDNA, the highest concentration is 350 ng/μL, followed by three 5-fold dilutions. Red asterisk indicates the band that was analyzed by Edman degradation. See **Figure S4** for analysis of DNA binding and sequence-specificity for cleavage activation. (C) Results of Edman degradation of CapH C-terminal cleavage product (red asterisk from panel (B), showing cleavage at residue R83. See **Figure S5** for full data. (D) Cartoon view of the CapH^67-107^ dimer, with R83 colored white and shown as sticks. (E) In vitro cleavage of MBP-CapH-GFP (wild-type or R83A mutant) by CapP, in the presence of 10 μM ssDNA. (F) GFP reporter assay showing effect of a CapP E98Q catalytic-dead mutant, CapH R83A mutant, or removal of CapH residues 83-107 (*capH* 1-82) on GFP expression. RNAP: RNA polymerase loading control. See full blot in **Figure S1A**. (G) CBASS expression reporter system with FLAG-NucC, showing effect of a CapH R83A mutant, or removal of CapH residues 83-107 (*capH* 1-82) on FLAG-NucC expression. α-RNAP: anti-RNA polymerase loading control. See full blot in **Figure S1B**.

Since we observed CapP-mediated CapH cleavage in cells, but not with purified proteins, we theorized that a soluble ligand present in cells is responsible for activating CapP. To test this idea, we performed in vitro cleavage assays with purified CapP and MBP-CapH-GFP in the presence of *E. coli* cell lysate. We observed robust activation of CapP in the presence of *E. coli* lysate, and stronger activation after the lysate was boiled and centrifuged to remove all proteins (**Figure S4B**). By fractionating boiled cell lysate using anion-exchange and size-exclusion chromatography, we found that CapP was most likely activated by a large, negatively-charged macromolecule (not shown). After observing a decrease in CapP activation upon treatment of the boiled cell lysate with DNase (not shown), we tested whether DNA could directly stimulate CapP. We found that single-stranded DNA strongly activates CapP (**Figure 5B**), while double-stranded DNA weakly activates CapP (**Figure 5B**), and single-stranded RNA does not stimulate CapP (**Figure S4C**). We found that single-stranded DNA as short as 5 bases long stimulated CapP, and that pyrimidines – particularly thymine – have the strongest stimulatory effect (**Figure S4C**). Finally, we found that CapP directly binds a single-stranded DNA oligonucleotide in vitro, but not an equivalent-length double-stranded DNA (**Figure S4D**). Further, CapP strongly binds poly-T DNA, weakly binds poly-C, and does not bind poly-A (**Figure S4E**). These data support a model in which CapP’s peptidase activity is stimulated by the binding of single-stranded DNA, particularly T-rich DNA.

We isolated the C-terminal product from CapP-mediated cleavage of MBP-CapH-GFP, and subjected it to Edman degradation to map the CapP cleavage site (**Figure 5C, S5**). We identified the cleavage site as between Phe82 and Arg83, within the first α-helix of the CapH C-terminal dimerization domain (**Figure 5C-D**). Confirming this finding, we found that a CapH Arg83 to alanine (R83A) mutant is not cleaved by CapP in vitro (**Figure 5E**). Finally, we found that in cells, truncation of CapH at residue 82 – mimicking CapP-cleaved CapH – resulted in strong expression as measured by both our GFP reporter system (**Figure 5F**) and our FLAG-NucC reporter (**Figure 5G**).

### CapP is activated by DNA damage

Our data showing that CapP is directly stimulated by single-stranded DNA in vitro suggests that in cells, it is activated by DNA damage. To test this idea, we coexpressed CapP and MBP-CapH-GFP in *E. coli* in the presence or absence of DNA damaging drugs. We found that in the presence of zeocin, a drug that induces DNA double-strand breaks (Chankova *et al*, 2007), CapP-mediated CapH cleavage was stimulated (**Figure 6A**). Using our FLAG-NucC reporter system, we found that both zeocin and mitomycin C, another damaging agent that causes DNA double-strand breaks through a separate mechanism (Kidane *et al*, 2004; Ayora *et al*, 2011), strongly stimulate CBASS expression within 60-120 minutes of adding the drugs (**Figure 6B-C**). This boost in CBASS expression was not observed in cells lacking *capP* (**Figure 6B-C**). Thus, CapH and CapP mediate increased CBASS expression upon DNA damage, even in the absence of phage infection.

**Figure 6.**
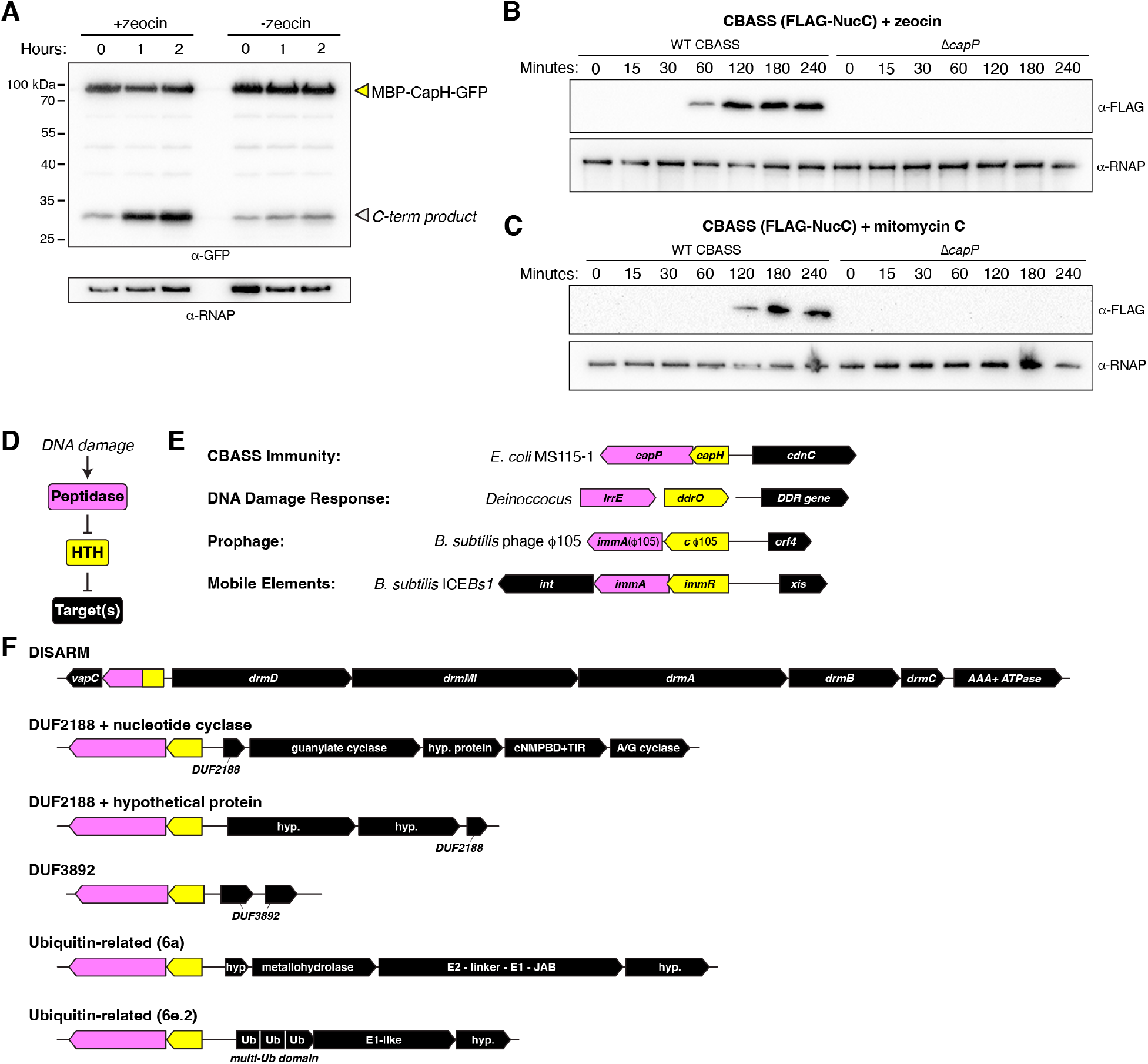
CapH and CapP induce CBASS expression in response to DNA damage. (A) Anti-GFP western blot showing coexpression in *E. coli* of an MBP-CapH-GFP fusion construct with wild-type CapP after exposure to zeocin (100 μg/mL). Full-length MBP-CapH-GFP is indicated with a yellow arrowhead, and the C-terminal product of CapP cleavage is indicated with a white arrowhead. α-RNAP: anti-RNA polymerase loading control. (B) Western blot of the CBASS expression reporter system with FLAG-NucC, showing FLAG-NucC expression after exposure to zeocin (100 μg/mL). α-RNAP: anti-RNA polymerase loading control. (C) Western blot of the CBASS expression reporter system with FLAG-NucC, showing FLAG-NucC expression after exposure to mitomycin C (1 μg/mL). α-RNAP: anti-RNA polymerase loading control. (D) Proposed signaling pathway for DNA damage-responsive transcriptional control systems in bacteria. (E) Diverse systems in bacteria that include an HTH transcriptional repressor (yellow) and a DNA damage-activated Zn^2+^ metallopeptidase (pink) that targets the transcriptional repressor for cleavage. (F) Known or likely bacterial defense systems associated with CapH (yellow) and CapP (pink)-like genes. The noted DISARM system from *Nocardia wallacei* FMUON74 encodes a *vapC*-like gene (NCBI accession # WP_187684394.1), a CapH-CapP fusion-like protein (WP_197986914.1), *drmD* (WP_187684395.1), drmMI (WP_187684396.1), drmA (WP_187684397.1), drmB ((WP_187684398.1), drmC (WP_187684399.1), and a AAA+ ATPase (WP_232110603.1). The noted DUF2188 + nucleotide cyclase system from *Tsuneonella flava* MS1-4 encodes a CapP-like protein (WP_102155989.1), a CapH-like protein (WP_102155988.1), a DUF2188 protein (WP_007013875.1), a predicted guanylate cyclase (WP_102155986.1), a hypothetical protein (WP_123639961.1), a predicted cyclic nucleotide monophosphate binding domain (cNMPBD) plus TIR domain protein (WP_102155984.1), and a predicted adenylate/guanylate cyclase (WP_102155983.1). The noted DUF2188 + hypothetical protein system from *Extensimonas perlucida* HX2-24 encodes a CapP-like protein (WP_144728453.1), a CapH-like protein (WP_144728455.1), two hypothetical proteins (WP_144728457.1 and WP_144728459.1), and a DUF2188 protein (WP_144729358.1). The noted DUF3892 system from *Sulfitobacter* sp. CW3 encodes a CapP-like protein (WP_037275352.1), a CapH-like protein (WP_037275354.1), and two DUF3892 proteins (WP_037275364.1 and WP_037275356.1). The noted ubiquitin-related (6a) system from *Methylobacterium oxalidis* NBRC 107715 encodes a CapP-like protein (WP_147028642.1), a CapH-like protein (WP_147028643.1), a hypothetical protein (WP_147028644.1), a predicted metallohydrolase (WP_147028653.1), a protein with predicted E2, E1, and JAB domains (WP_147028645.1), and a second hypothetical protein (WP_147028646.1). The noted ubiquitin-related (6e.2) system from *Mixta intestinalis* SRCM103226 encodes a CapP-like protein (WP_160622475.1), a CapH-like protein (WP_048227226.1), a predicted multi-ubiquitin domain protein (WP_053069300.1), a predicted E1-like protein (WP_160622476.1), and a hypothetical protein (WP_18149987.1).

### CapH and CapP are members of a broadly conserved family of DNA damage response proteins

CapH and CapP show strong structural and functional similarity to the *Deinococcus* proteins DdrO and IrrE, with both systems inducing expression of target genes upon DNA damage through metallopeptidase cleavage of a transcriptional repressor (**Figure 6D-E**). DNA damage is a universal signal of cell stress, and as such is a major signal to induce lysogenic phage (prophage) to a switch to the lytic life cycle, and to induce mobility of integrative and conjugative elements (ICE elements) (Bæk *et al*, 2003; Auchtung *et al*, 2005). Strikingly, the use of an HTH family transcriptional repressor coupled with a DNA damage-stimulated metallopeptidase is shared in some prophages and ICE elements. For example, *Bacillus subtilis* mobile element ICE*Bs*1 and prophage ɸ105 each encode an HTH-family transcriptional repressor (ImmR and cɸ105, respectively) that strongly represses the expression of genes responsible for excision of these elements from the genome, and a Zn^2+^ metallopeptidase (ImmA) that cleaves the HTH protein upon DNA damage to relieve repression and induce excision (**Figure 6E**) (Bose *et al*, 2008; Bose & Grossman, 2011).

The structural and functional parallels between CapH/CapP, DdrO/IrrE, and ImmR/ImmA suggest that these proteins represent a broadly conserved family of DNA damage responsive transcriptional regulators. We used the FlaGs (<u>Fla</u>nking <u>G</u>ene<u>s</u>) server to search for other instances of CapH/CapP-like proteins and identify their associated operons (Saha *et al*, 2021). We identified a broad range of operons associated with *capH* and *capP*-like genes, with all of them sharing the location of *capH* and *capP* upstream of, and oriented on the opposite strand as, the associated operon (**Figure 6F**). Most of these systems represent known or putative defense systems, notably including a set of DISARM anti-phage systems associated with a CapH-CapP fusion protein. We identified three sets of operons encoding proteins of the DUF2188 or DUF3892 families, which are uncharacterized but have been previously linked to bacterial defense (Burroughs & Aravind, 2020). Both DUF2188 and DUF3892 proteins have also been identified in operons containing HTH and metallopeptidase genes, paralleling our findings (Burroughs & Aravind, 2020). We also identified operons encoding proteins related to eukaryotic ubiquitin signaling machinery, including so-called 6a systems that encode a large protein with E2-like, E1-like, and JAB protease-like domains (Burroughs *et al*, 2009; Iyer *et al*, 2006). Notably, this protein shares strong homology to the Cap2 protein in Type II CBASS systems (also classified as 6b systems), which conjugates the C-terminus of its cognate CD-NTase to an unknown target to regulate anti-phage signaling (Ledvina *et al*, 2022). We also identified CapH and CapP-like proteins associated with 6e systems, which encode a protein predicted to possess multiple ubiquitin-like β-grasp domains and an E1-like protein (**Figure 6F**). Thus, CapH and CapP-like regulators are associated with a broad range of bacterial signaling pathways with known or predicted roles in anti-phage or stress responses.

## Discussion

Here, we identify a pair of proteins – CapH and CapP – that are associated with hundreds of CBASS anti-phage systems, and show that they function together to regulate CBASS expression. In the basal state, the helix-turn-helix protein CapH forms oligomers and binds the promoter region of its associated CBASS system to repress transcription. With its distinctive cysteine switch motif, CapP is maintained in an inactive state in this mode. In *E. coli* MS115-1 CBASS, the resulting low-level basal expression of the CBASS core genes, which is undetectable by western blotting in our reporter system, is apparently sufficient to provide protection against bacteriophage λ infection. Despite the extremely low levels of the core CBASS proteins in this repressed state, phage infection is sensed by the system’s HORMA domain proteins, activating CdnC to produce a cyclic tri-AMP second messenger. Cyclic tri-AMP in turn activates NucC, which destroys the host genome to kill the cell and abort the infection (**Figure 7A**) (Ye *et al*, 2020; Lau *et al*, 2020).

**Figure 7.**
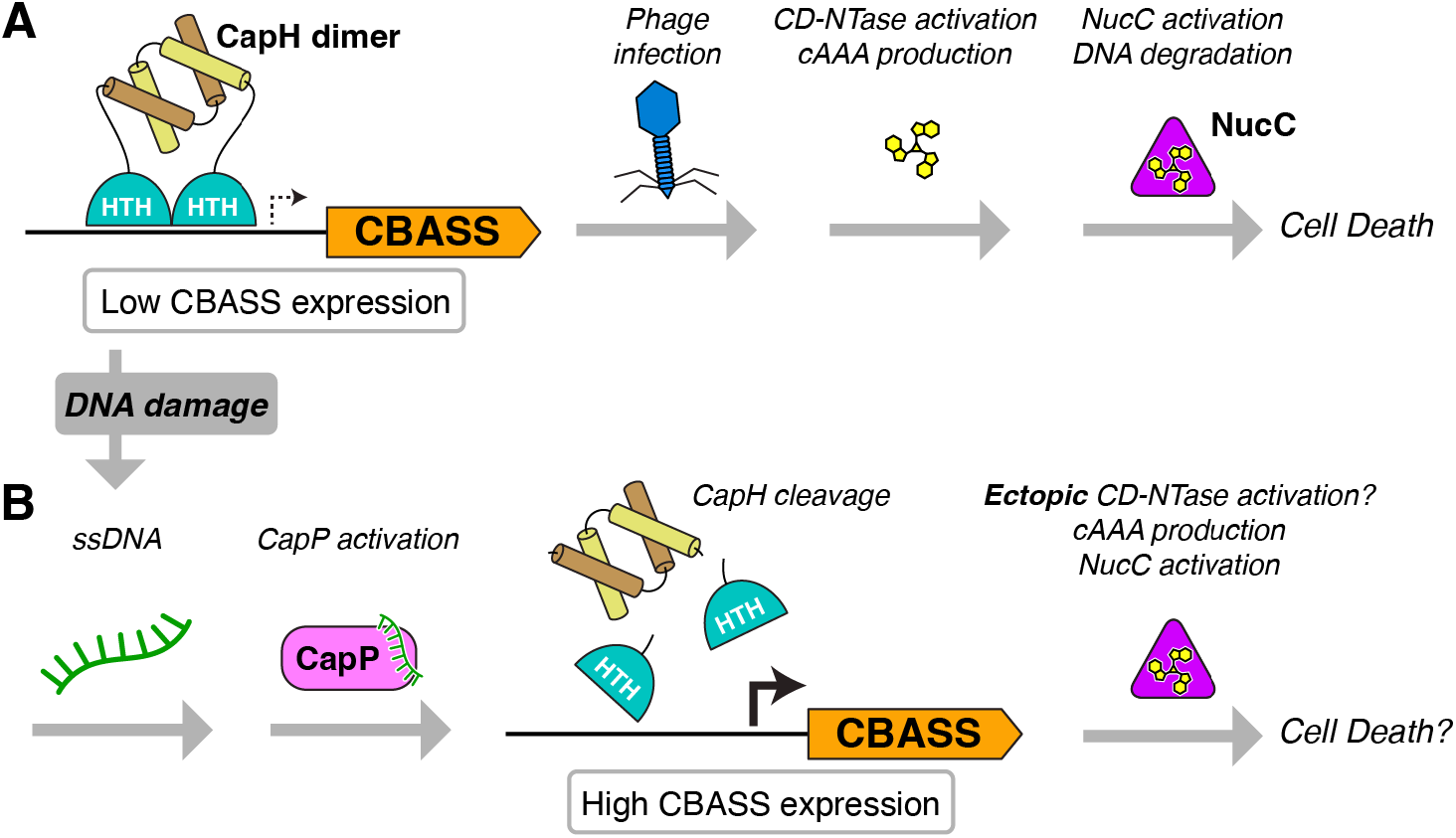
Model for CapH/CapP function in CBASS-mediated immunity. (A) In the basal state, CapH dimers (depicted) or monomers (not shown) bind their cognate CBASS promoter and repress transcription. Low-level CBASS expression is sufficient for detection of a lytic phage infection, CD-NTase activation and second messenger production (in *E. coli* MS115-1 CBASS, cyclic tri-AMP/cAAA), followed by effector activation (in *E. coli* MS115-1, NucC) and cell death. (B) Upon DNA damage induced by lysogenic phage infection, activation of other anti-phage immune systems, or environmental stimuli, CapP is activated and cleaves CapH. CapH cleavage and release from the CBASS promoter mediates a dramatic increase in CBASS activation, resulting in ectopic CD-NTase activation, second messenger production, effector activation, and cell death.

If CapH and CapP are not required for protection against phage infection, what is their role? We find that DNA damage strongly activates CapP-mediated CapH cleavage through the production of T-rich single-stranded DNA, and that this activation results in a dramatic increase in CBASS expression. We propose that this pathway represents a second path for CBASS activation that directly responds to DNA damage rather than phage infection. DNA damage may arise from multiple sources, including prophage induction (Murphy, 1998; Datsenko & Wanner, 2000) or the action of other defense systems like restriction/modification or CRISPR/Cas systems. We propose that the dramatic increase in CBASS expression upon DNA damage may mediate ectopic activation of the system’s CD-NTase, bypassing the canonical activation mechanism to activate the system’s effector protein and kill the cell (**Figure 7B**). In this manner, CBASS systems encoding CapH and CapP could sense and respond to a broader variety of stress stimuli than systems lacking these regulators, potentially enabling them to cooperate with other immune systems like restriction/modification and CRISPR/Cas systems.

The mechanism of CapH/CapP-mediated transcriptional regulation parallels that of the *Deinococcus* DNA damage response proteins DdrO/IrrE (Devigne *et al*, 2015; de Groot *et al*, 2019; Ludanyi *et al*, 2014), and of the ImmR/ImmA proteins that induce excision of prophages and ICE elements in response to DNA damage. While ImmA responds to DNA damage through association with RecA in a SOS response-dependent manner, or through the signaling regulator RapI (Bose *et al*, 2008; Bose & Grossman, 2011), how IrrE is activated remains unknown. Given that both CapP and IrrE share a C-terminal GAF domain, which in other proteins interacts with cyclic nucleotides (cAMP or cGMP), suggests that IrrE may share a similar mechanism of ssDNA-stimulated peptidase activity (Gross-Langenhoff *et al*, 2006; Martinez *et al*, 2008; Ho *et al*, 2000).

In addition to IrrE/DdrO and ImmR/ImmA, we identified a broad set of known or putative defense systems associated with CapH and CapP-like regulators. These include a subset of DISARM systems, which function similarly to restriction/modification systems in phage defense (Ofir *et al*, 2018). CapP and CapH also appear in operons encoding DUF2188 and DUF3892 proteins, which have both been predicted to participate in anti-phage defense (Burroughs & Aravind, 2020). Finally, the association of CapH/CapP with uncharacterized operons encoding proteins related to ubiquitin signaling machinery suggests that these operons, too, may play a role in defense against phage infection or other stresses.

Our identification of a mechanism enabling a single defense system to respond to multiple stimuli parallels the recent discovery and characterization of BrxR/CapW transcriptional regulators, which are associated with a variety of immune systems including BREX and CBASS anti-phage systems (Picton *et al*, 2021; Blankenchip *et al*, 2022; Luyten *et al*, 2022). CapW drives increased expression of its associated CBASS systems upon phage infection, but as with CapH/CapP-associated systems, this increased expression is not required for protection against lytic phage (Blankenchip *et al*, 2022). Similarly, BrxR controls expression of its associated BREX systems, but is not required for anti-phage immunity (Luyten *et al*, 2022). While the activating signal of BrxR/CapW is not known, these data suggest that the protein controls activation of CBASS and BREX systems in response to signals other than phage infection. As with CapH and CapP, BrxR/CapW may enable their associated defense systems to act as a second line of defense in coordination with restriction/modification or CRISPR/Cas systems (Luyten *et al*, 2022). More broadly, there may exist a range of signaling mechanisms that enable crosstalk between distinct defense systems in bacteria, mediating these systems’ cooperation and integration into a comprehensive, multi-faceted immune system.

## Materials and Methods

### Bioinformatics

To comprehensively search CBASS systems, we exported the genomic DNA sequences within 10 kb of 6233 previously-reported CD-NTases (Cohen *et al*, 2019) using the Integrated Microbial Genomes (IMG) database at the DOE Joint Genome Institute (https://img.jgi.doe.gov). We used NCBI Genome Workbench (https://www.ncbi.nlm.nih.gov/tools/gbench/) to perform custom TBLASTN searches for proteins related to *E. coli* MS115-1 CapP (NCBI sequence ID WP_001290439.1; **Table S1**). CBASS system type and effector assignments for each hit were taken from Cohen et al. (Cohen *et al*, 2019) and manually updated. Each hit was manually inspected for the presence of CapH and CapP.

### Cloning, expression, and protein purification

All proteins were cloned into UC Berkeley Macrolab vector 2BT (Addgene #29666; encoding an N-terminal TEV protease-cleavable His_6_-tag). Proteins used were: *E. coli* MS115-1 CapH (NCBI sequence ID WP_001515173.1), *E. coli* MS115-1 CapP (NCBI sequence ID WP_001290439.1), and *Thauera* sp. K11 CapP (NCBI sequence ID WP_096453114.1). Proteins were expressed in *E. coli* strain Rosetta 2 (DE3) pLysS (EMD Millipore, Billerica MA). Cultures were grown at 37°C to A_600_=0.5, then induced with 0.25 mM IPTG and shifted to 20°C for 15 hours. Cells were harvested by centrifugation and resuspended in buffer A (20 mM Tris pH 7.5, 10% glycerol) plus 300 mM NaCl, 10 mM imidazole, and 5 mM β-mercaptoethanol (10 µM ZnCl_2_ was added to buffers for CapP). Proteins were purified by Ni^2+^-affinity (Ni-NTA agarose, Qiagen) then passed over an anion-exchange column (Hitrap Q HP, Cytiva) in Buffer A plus 5 mM β-mercaptoethanol and 0.1-1 M NaCl, collecting flow-through or peak fractions. Tags were cleaved with TEV protease (Tropea *et al*, 2009), and cleaved protein was passed over another Ni^2+^ column (collecting flow-through fractions) to remove uncleaved protein, cleaved tags, and tagged TEV protease. The protein was passed over a size exclusion column (Superdex 200, Cytiva) in buffer GF (buffer A plus 300 mM NaCl and 1 mM dithiothreitol (DTT)), then concentrated by ultrafiltration (Amicon Ultra, EMD Millipore) to 10-20 mg/ml and stored at 4°C. For selenomethionine derivatization, protein expression was carried out in M9 minimal media supplemented with amino acids plus selenomethionine prior to IPTG induction (Van Duyne *et al*, 1993), and proteins were exchanged into buffer containing 1 mM tris(2-carboxyethyl)phosphine (TCEP) after purification to maintain the selenomethionine residues in the reduced state.

### Crystallization and structure determination

For crystallization of *E. coli* MS115-1 CapH^NTD^ (residues 2-67), protein was concentrated to 18 mg/mL in crystallization buffer (20 mM Tris-HCl pH 8.5, 150 mM NaCl, 1 mM DTT) then mixed 1:1 with well solution containing 0.1 M Ammonium acetate pH 4.5 and 2 M Ammonium sulfate in hanging-drop format. Crystals were cryoprotected with an additional 25% sucrose and flash-frozen in liquid nitrogen. Diffraction data to 1.02 Å resolution were collected at Advanced Light Source beamline 5.0.2 (see support statement below) and processed with the DIALS data-processing pipeline (https://dials.github.io) (Beilsten-Edmands *et al*, 2020). We determined the structure by molecular replacement in PHASER (McCoy *et al*, 2007), using an ideal alpha-helix as a search model. We manually rebuilt the initial model in COOT (Emsley *et al*, 2010), and refined in phenix.refine (Afonine *et al*, 2012) using individual positional and anisotropic B-factor refinement for non-hydrogen atoms, and riding hydrogens (**Table S2**).

For crystallization of *E. coli* MS115-1 CapH^CTD^(I99M) (residues 67-107 with Ile99 to Met mutation), protein was concentrated to 8 mg/mL in crystallization buffer (20 mM Tris-HCl pH 8.5, 150 mM NaCl, 1 mM DTT) then mixed 1:1 with well solution containing 0.1 M HEPES pH 7.5, 25 mM MgCl_2_, and 30% PEG 550 monomethyl ether in hanging-drop format. Crystals were cryoprotected with an additional 13% PEG 550 monomethyl ether and 10% glycerol, and flash-frozen in liquid nitrogen. Diffraction data to 1.26 Å resolution were collected at Advanced Photon Source beamline 24ID-C (see support statement below) and processed with the RAPD data-processing pipeline, which uses XDS (Kabsch, 2010)for data indexing and reduction, AIMLESS (Evans & Murshudov, 2013) for scaling, and TRUNCATE (French & Wilson, 1978) for conversion to structure factors. We determined the structure by molecular replacement in PHASER (McCoy *et al*, 2007), using an ideal α-helix as a search model. We manually rebuilt the initial model in COOT (Emsley *et al*, 2010), and refined in phenix.refine (Afonine *et al*, 2012) using individual positional and anisotropic B-factor refinement for non-hydrogen atoms, and riding hydrogens (**Table S2**).

For crystallization of *E. coli* MS115-1 CapH^CTD^ (residues 67-107), protein was concentrated to 21 mg/mL in crystallization buffer (20 mM Tris-HCl pH 8.5, 150 mM NaCl, 1 mM DTT) then mixed 1:1 with well solution containing 0.1 M sodium citrate pH 3.0 and 1.6 M lithium sulfate in hanging-drop format. Crystals were cryoprotected with an additional 20% ethylene glycol and flash-frozen in liquid nitrogen. Diffraction data to 1.75 Å resolution were collected at Advanced Light Source beamline 5.0.2 (see support statement below) and processed with the DIALS data-processing pipeline (https://dials.github.io) (Beilsten-Edmands *et al*, 2020).We determined the structure by molecular replacement in PHASER (McCoy *et al*, 2007), using the structure of CapH^CTD^(I99M) as a search model. We manually rebuilt the initial model in COOT (Emsley *et al*, 2010), and refined in phenix.refine (Afonine *et al*, 2012) using individual positional and B-factor refinement, and riding hydrogens (**Table S2**).

For crystallization of *Thauera* sp. K11 CapP, protein was concentrated to 9 mg/mL in crystallization buffer (20 mM Tris-HCl pH 7.5, 100 mM NaCl, 1 mM DTT) then mixed 1:1 with well solution containing 0.1 M CHES pH 9.5, 0.3 M NaCl, and 1.8 M lithium sulfate in hanging-drop format. Crystals were cryoprotected with an additional 30% glycerol and flash-frozen in liquid nitrogen. Diffraction data for both native and selenomethionine-derivatized crystals were collected at the Stanford Synchrotron Radiation Lightsource beamline 9-2 (see support statement below) and processed with the autoxds data-processing pipeline, which uses XDS (Kabsch, 2010) for data indexing and reduction, AIMLESS (Evans & Murshudov, 2013) for scaling, and TRUNCATE (French & Wilson, 1978) for conversion to structure factors.). We determined the structure by single-wavelength anomalous diffraction (SAD) methods in PHASER (McCoy *et al*, 2007) using a 1.6 Å resolution dataset from selenomethionine-derivatized protein. We manually rebuilt the initial model in COOT (Emsley *et al*, 2010), and refined against a 1.35 Å resolution native dataset in phenix.refine (Afonine *et al*, 2012) using individual positional and anisotropic B-factor refinement, and riding hydrogens (**Table S2**).

### Beamline Support Statements

#### ALS beamline 5.0.2

The Berkeley Center for Structural Biology is supported in part by the Howard Hughes Medical Institute. The Advanced Light Source is a Department of Energy Office of Science User Facility under Contract No. DE-AC02-05CH11231. The Pilatus detector on 5.0.1. was funded under NIH grant S10OD021832. The ALS-ENABLE beamlines are supported in part by the National Institutes of Health, National Institute of General Medical Sciences, grant P30 GM124169.

#### APS beamline 24ID-C

This work is based upon research conducted at the Northeastern Collaborative Access Team beamlines, which are funded by the National Institute of General Medical Sciences from the National Institutes of Health (P30 GM124165). This research used resources of the Advanced Photon Source, a U.S. Department of Energy (DOE) Office of Science User Facility operated for the DOE Office of Science by Argonne National Laboratory under Contract No. DE-AC02-06CH11357.

#### SSRL beamline 9-2

Use of the Stanford Synchrotron Radiation Lightsource, SLAC National Accelerator Laboratory, is supported by the U.S. Department of Energy, Office of Science, Office of Basic Energy Sciences under Contract No. DE-AC02-76SF00515. The SSRL Structural Molecular Biology Program is supported by the DOE Office of Biological and Environmental Research, and by the National Institutes of Health, National Institute of General Medical Sciences (P30GM133894). The contents of this publication are solely the responsibility of the authors and do not necessarily represent the official views of NIGMS or NIH.

### SEC-MALS

For characterization of oligomeric state by size exclusion chromatography coupled to multi-angle light scattering (SEC-MALS), 100 μL of purified protein/complex at 2-5 mg/mL was injected onto a Superdex 200 Increase 10/300 GL column (Cytiva) in a buffer containing 20 mM HEPES pH 8.5, 300 mM NaCl, 5% glycerol, and 1 mM DTT. Light scattering and refractive index profiles were collected by miniDAWN TREOS and Optilab T-rEX detectors (Wyatt Technology), respectively, and molecular weight was calculated using ASTRA v. 8 software (Wyatt Technology).

### DNA-binding assays

For characterization of DNA binding by fluorescence polarization assays, a 22 bp double-stranded DNA was produced by annealing complementary oligos, one of which was 5’-6-FAM labeled (the same oligonucleotide was used without annealing for ssDNA binding studies). Binding reactions (30 µL) contained 25 mM Tris pH 7.5, 50 mM NaCl, 5% glycerol, 5 mM MgCl2, 1 mM DTT, 0.01% nonidet p40 substitute, 50 nM DNA, and the indicated amounts of protein. After a 10 minute incubation at room temperature, fluorescence polarization was read using a Tecan Infinite M1000 PRO fluorescence plate reader, and binding data were analyzed with Graphpad Prism v.9.2.0 using a single-site binding model.

### CapP cleavage assays

For detection of CapP activity in cells, *E. coli* MS115-1 CapP was co-expressed with a construct of *E. coli* MS115-1 CapH fused to an N-terminal His_6_-maltose binding protein (MBP) tag and a C-terminal green fluorescent protein (GFP) tag (MBP-CapH-GFP) in *E. coli* Rosetta 2 (DE3) pLysS cells at 37°C. Protein expression was induced with 0.25 mM IPTG for 2-4 hours, then samples were removed for analysis by western blot (see below). For CapP activity in cells in response to zeocin, *E. coli* Rosetta 2 (DE3) pLysS cells were grown at 37°C until reaching an OD_600_ of 0.3. Protein expression was induced with 0.25 mM IPTG for 1 hour, then zeocin was added at a concentration of 100 μg/mL. 500 μL of sample was taken at 1 hour and 2 hours post-antibiotic and centrifuged 10,000xg for 1 minute. Pelleted cells were resuspended in 50 uL of 2xSDS sample buffer and analyzed by western blot (see below).

For detection of CapP activity using purified proteins, 20 μL reactions containing 10 μM purified CapP and 10 μM purified MBP-CapH-GFP in a buffer containing 50 mM Tris pH 7.5, 300 mM NaCl and 5 μM ZnCl_2_ were incubated at 37°C for 2 hours, then added to 20 uL 2xSDS sample buffer. 10 μL of each sample was loaded and separated by SDS-PAGE and Coomassie-stained for visualization. For reactions containing *E. coli* cell lysate, log-phage *E. coli* Rosetta 2 (DE3) pLysS cells were lysed by sonication in a minimum volume of buffer A, then centrifuged to remove cell debris. Optionally, the lysate was incubated in a boiling water bath for 10 minutes, then centrifuged again to remove denatured proteins. 10 μL of cell lysate was added to reactions with purified CapP and MBP-CapH-GFP. For reactions containing DNA, nucleic acid was added at a concentration of 350 μg/mL and subsequent serial 5-fold dilutions, then incubated in the buffer as above for 2 hours. For reactions containing DNA or RNA oligos, nucleic acid was added at a concentration of 10 μM, then incubated in the buffer as above for 2 hours.

### Edman degradation

For Edman degradation, MBP-CapH-GFP was treated with CapP plus single-stranded DNA, separated by SDS-PAGE, then transferred to a PVDF membrane and visualized with Coomassie staining. The band representing the C-terminal cleavage product of MBP-CapH-GFP was cut out and analyzed by Edman degradation at the UC Davis Molecular Structure Facility (http://msf.ucdavis.edu).

### Western Blots

For CapH repressor assays, cells containing plasmids with the MS115-1 system were grown in 5 mL of LB plus the selection marker at 37°C until they reached an OD_600_ of 0.3 – 0.5. Cultures were adjusted to an OD_600_ of 0.3, then an aliquot of 500 μL was removed and centrifuged for 1 minute at 10,000 x g to pellet the cells. The supernatant was removed and the cells were resuspended in 50 μL of 2xSDS sample buffer (125 mM Tris pH 6.8, 20% Glycerol, 4% SDS, 200 mM DTT, 180 μM bromophenol blue). 10μL of each sample was loaded onto a 12% SDS-PAGE gel for separation and transferred to a PVDF membrane using the Bio-Rad Trans-Blot Turbo Transfer System. Membranes were blocked for 1 hour in 5% milk in TBST (100 mM Tris pH 7.5, 150 mM NaCl, 0.1% Tween-20) at room temperature with shaking, then incubated with anti-FLAG mouse antibody (Sigma-Aldrich) or anti-GFP antibody (Roche) at 1:3,000 diluted in 5% milk in TBST for 1 hour at room temperature with shaking. Membranes were washed 3 times with 10 mL of TBST then incubated with anti-mouse goat antibody conjugated to horseradish peroxidase (Millipore Sigma) at 1:30,000 diluted in 5% milk in TBST for 1 hour at room temperature with shaking. Membranes were again washed 3 times with 10 mL TBST then incubated with Pierce ECL Plus substrate for 2 minutes and then imaged on a Bio-Rad ChemiDoc imager. Membranes were stripped with a solution of 200 mM glycine pH 2.2, 0.1% SDS, and 1% Tween 20 for 20 minutes at room temperature, then washed with TBST and re-blocked with 5% milk overnight at 4°C. Membranes were re-blotted using the same procedure as initial blot, but replacing primary antibody with anti-RNA polymerase B mouse antibody (clone NT63; BioLegend #10019-878).

### DNA damage assays

For antibiotic treatment time course western blots, *E. coli* JP313 containing reporter plasmids were grown at 37°C until reaching an OD_600_ of 0.1. Cultures were moved to 30°C for 10 minutes, then antibiotics were added at a concentration of 100 μg/mL for zeocin and 1 μg/mL for mitomycin C. At each timepoint, 1 mL of culture was removed and resuspended in 2xSDS sample buffer with the volume adjusted to an equal concentration of cells per sample. Western blots were completed as described above

### Bacteriophage infectivity

For plaque quantification assays, strains were grown from single colonies in 5 mL of LB with ampicillin (100 ug/mL) until reaching log phase (0.3-0.6). 500 uL of culture was transferred to a 5 mL conical tube to which 10 uL of 10-fold dilutions of lambda cI-in phage buffer () was added. Tubes were incubated at room temperature for 20 minutes after which 4.5 mL of 0.35% top agar was added mixed, then poured onto LB plates containing carbinocillin. Plates were incubated at 37°C for 16 hours, then plaques were counted.

For infection time course western blots, *E. coli* JP313 cells containing reporter plasmids were grown at 37°C until reaching an OD_600_ of 0.3. Cultures were moved to 30°C for 10 minutes, then λ cI-was added at an MOI of 10. At each timepoint, 500 μL of sample was removed and centrifuged 10,000xg for 1 minute. Cell pellets were resuspended in 50 μL 2xSDS sample buffer and analyzed by western blot (see above).

### Microscopy

For fluorescence microscopy imaging, each sample was grown as liquid culture at 30°C and induced with 0.2mM IPTG 30 minutes before of imaging. Cells were infected with Lambda CI- at MOI 2.5 as 0 MPI. Cells were harvested at required timepoints and were briefly centrifuged (3300 x g for 30 seconds). After resuspending the cells with 20 µl volume, 5 µl of the samples were transferred onto an agarose pad containing 1% agarose and 20% LB medium for microscopy and stained with 1 µg/mL FM4-64 and 2 mg/mL DAPI. Microscopy was performed using by a Deltavision Elite System (GE Healthcare).

## Acknowledgements

The authors thank John Schulze at the UC Davis Molecular Structure Facility for assistance with Edman Degradation, and members of the Corbett lab and Aaron Whiteley for helpful feedback. K.D.C. acknowledges support from the National Institutes of Health, R21 AI148814 and R35 GM144121. R.K.L. was supported by the UCSD Quantitative and Integrative Physiology Training Grant (NIH T32 GM127235) and an individual National Institutes of Health Predoctoral Fellowship (F31 GM137600).

## Supplemental Information

**Figure S1.**
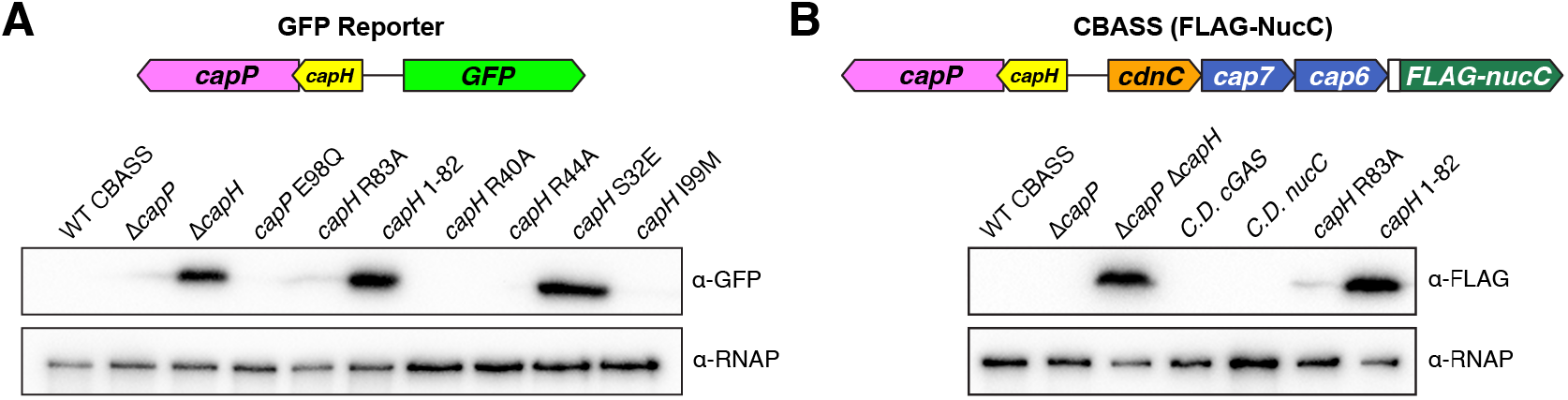
CBASS expression reporter systems. (A) Full western blot for GFP reporter assay shown in Figures 1D, **2E, 5F**. (B) Full western blot for FLAG-NucC expression reporter assay shown in Figures 1E**, 5G**.

**Figure S2.**
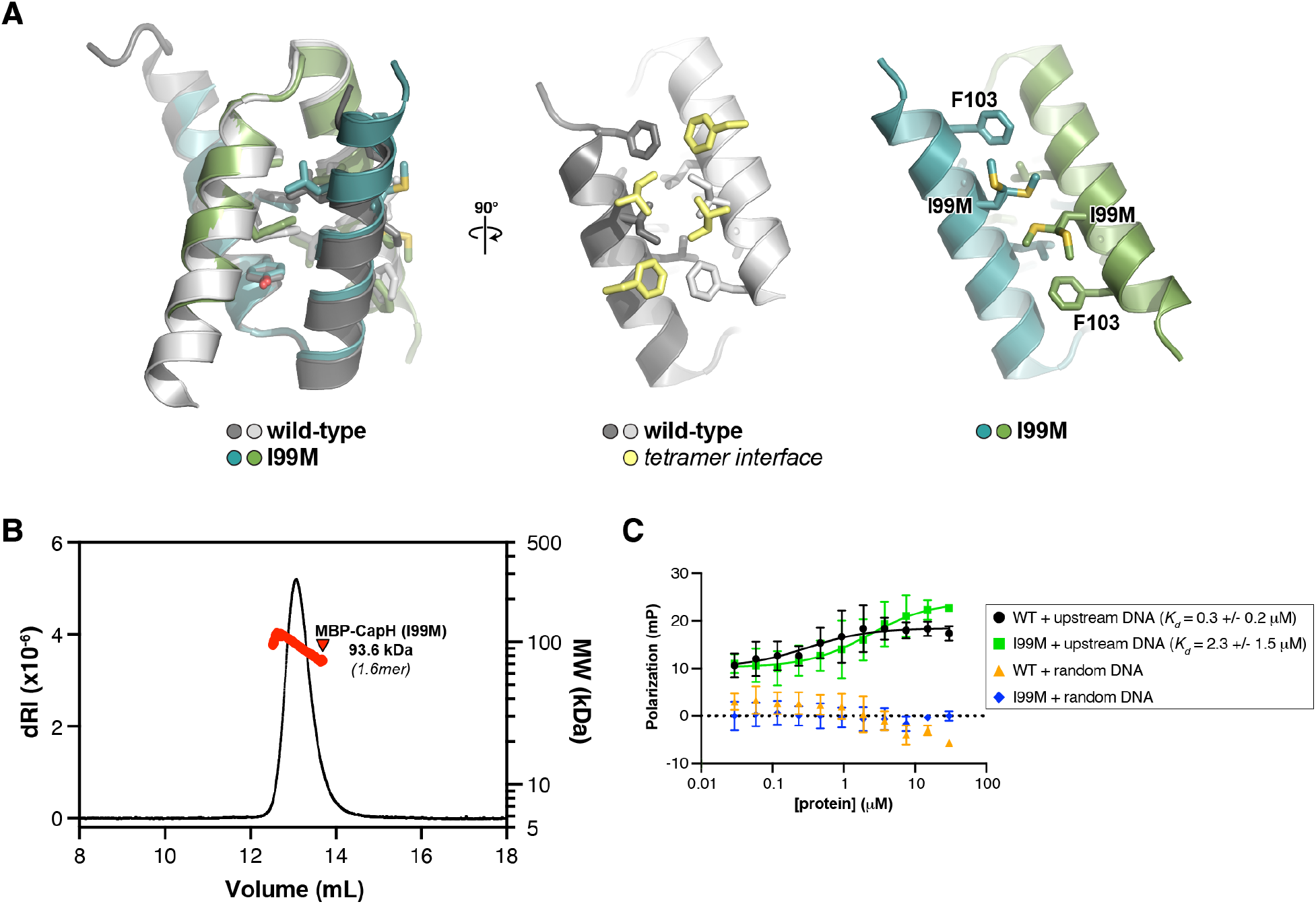
The CapH I99M mutation disrupts tetramer formation. (A) *Left:* Cartoon view of the *E. coli* MS115-1 CapH^CTD^ (I99M) structure, with the two protomers colored blue and green. Overlaid is the structure of wild-type CapH^CTD^ (dark gray/light gray). *Center:* Views of the wild-type CapH^CTD^ tetramerization interface, with the dimer shown in dark gray/light gray and interacting residues from the opposite dimer in light yellow. Right: View of the CapH^CTD^ (I99M) tetramerization interface, showing the position of the I99M mutation that disrupts tetramer formation. (B) SEC-MALS analysis of MBP-fused CapH (I99M) mutant (monomer molecular weight 57.0 kDa), showing that it forms a homodimer. (C) Fluorescence polarization DNA binding assay showing that the CapH I99M mutation reduces, but does not eliminate, the DNA binding ability of CapH.

**Figure S3.**
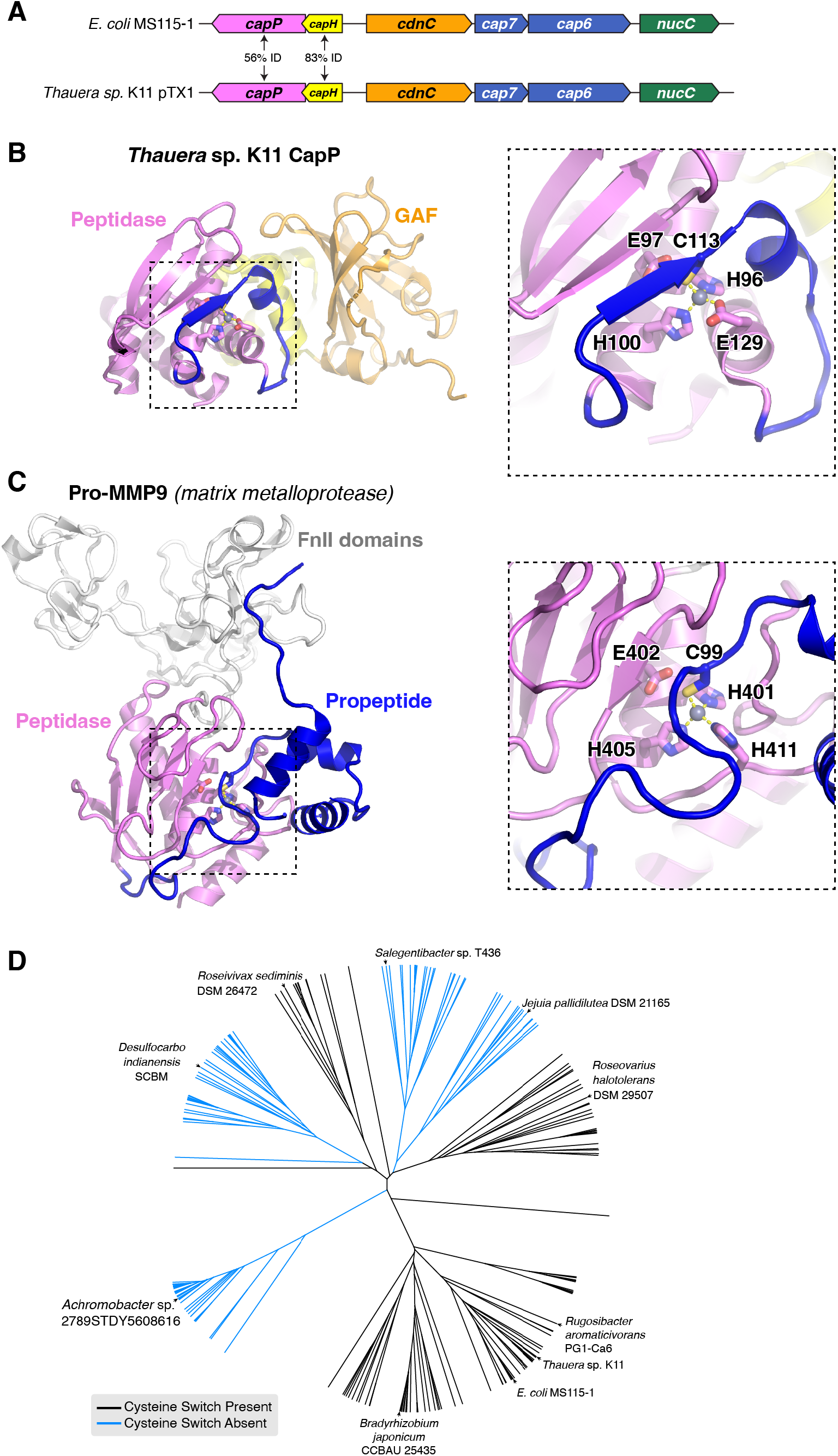
CBASS-associated CapP contains an internal cysteine switch. (A) Operon schematic of the *Thauera* sp. K11 CBASS system, compared to the *E. coli* MS115-1 system, with sequence identity between the two systems’ CapH and CapP proteins indicated. (B) Structure of *Thauera* sp. K11 CapP, with closeup of its Zn^2+^ metallopeptidase domain (pink) with internal cysteine switch loop (blue) and cysteine switch residue (C113). (C) Structure of human matrix metalloprotease MMP9 (PDB IF 1L6J; (Elkins *et al*, 2002)), with closeup of its Zn^2+^ metallopeptidase domain (pink) and N-terminal cysteine switch domain (blue) and cysteine switch residue (C99). The orientation of the Zn2+ metallopeptidase domains in panels B and C is identical. (D) Evolutionary tree of 408 CBASS-associated CapP proteins, colored by the presence (black) or absence (blue) of the internal cysteine switch.

**Figure S4.**
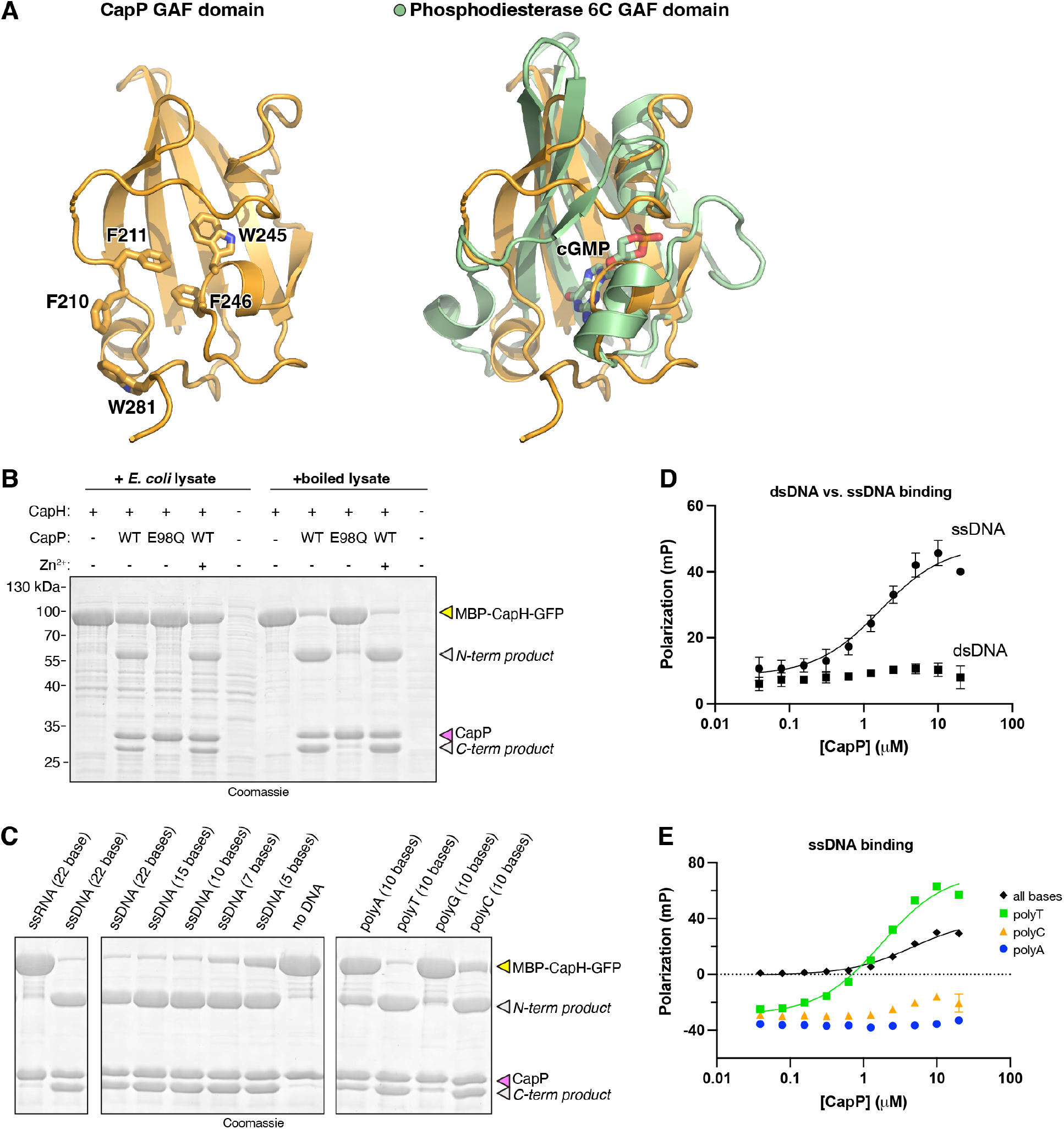
Structure and DNA-mediated activation of CapP. (A) Structural overlay between the *Thauera* sp. K11 CapP GAF domain (orange) and the GAF domain of phosphodiesterase 6C bound to cyclic GMP (green; PDB ID 3DBA; (Martinez *et al*, 2008)). Surface-exposed aromatic residues potentially involved in ligand binding are shown as sticks in the left panel. (B) In vitro cleavage assay with purified *E. coli* MS115-1 CapP (wild-type or catalytic-dead E98Q mutant), MBP-CapH-GFP, and *E. coli* cell lysate. (C) In vitro cleavage assay with purified *E. coli* MS115-1 CapP (wild-type or catalytic-dead E98Q mutant), MBP-CapH-GFP, and the indicated nucleic acids. (D) Fluorescence polarization DNA binding assay for *E. coli* MS115-1 CapP and either single-stranded DNA (circles and solid line; *K_d_*=1.7 +/- 0.6 μM) or double-stranded DNA (squares; no binding detected). (E) Fluorescence polarization DNA binding assay for *E. coli* MS115-1 CapP and single-stranded DNAs including a random sequence with all four bases (black diamonds; *K_d_*=4.7 +/- 1.2 μM), poly-T (green squares; *K_d_*=1.8 +/- 0.3 μM), poly-C (orange triangles; no binding detected), or poly-A (blue circles; no binding detected).

**Figure S5.**
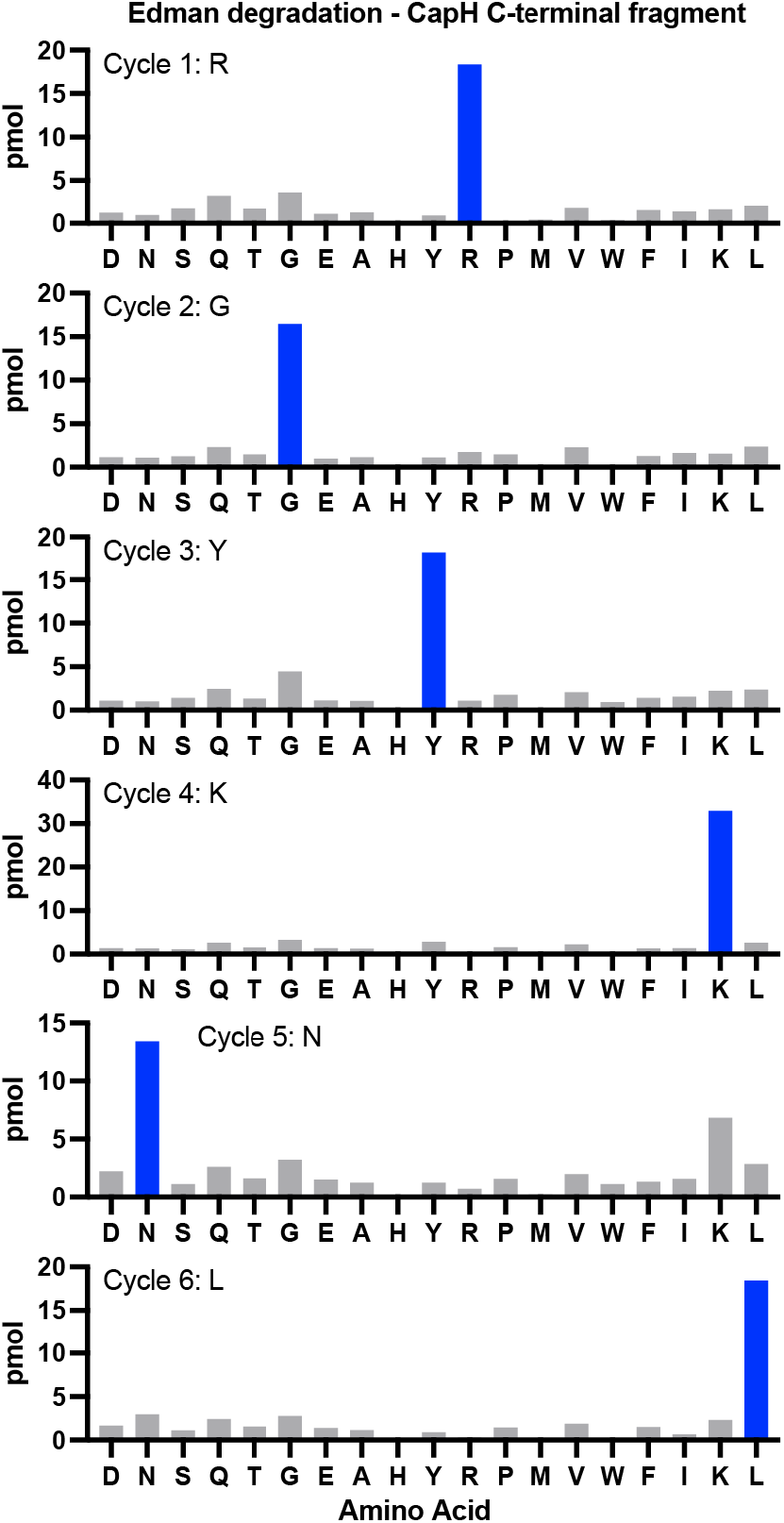
Edman degradation of CapP cleavage product. Bar graphs showing picomoles (pmol) of each amino acid detected by Edman degradation of the CapP-mediated cleavage product MBP-CapH-GFP (denoted by red asterisk in Figure 5B). The dominant amino acid for each cycle is shown as a blue bar. The inferred N-terminal sequence (RGYKNL) matches CapH residues 83-88 (see Figure 5C).

**Table S1.** CapH and CapP regulators in CBASS systems. *(see attached Microsoft Excel workbook)*

**Table S2.** Crystallographic data and refinement.

|  | *Thauera* sp. K11<br>CapP SeMet | *Thauera* sp. K11<br>CapP Native | *E. coli* MS115-1<br>CapH NTD | *E. coli* MS115-1<br>CapH CTD | *E. coli* MS115-1<br>CapH CTD (I99M) |
| --- | --- | --- | --- | --- | --- |
| **Data collection** |  |  |  |  |  |
| Synchrotron/Beamline | SSRL 9-2 | SSRL 9-2 | ALS 5.0.2 | ALS 5.0.2 | APS 24ID-C |
| Date collected | 02/14/2020 | 02/14/2020 | 04/25/2021 | 5/14/21 | 07/11/21 |
| Resolution (Å) | 40-1.6 | 40-1.35 | 40-1.02 | 100-1.75 | 59-1.26 |
| Wavelength (Å) | 0.97892 | 0.97892 | 1.00003 | 1.00004 | 0.97918 |
| Space Group | P4<sub>2</sub>2<sub>1</sub>2 | P4<sub>2</sub>2<sub>1</sub>2 | P2<sub>1</sub>2<sub>1</sub>2<sub>1</sub> | P2<sub>1</sub> | P4<sub>3</sub>2<sub>1</sub>2 |
| Unit Cell Dimensions (a, b, c) Å | 95.28, 95.28,<br>103.95 | 95.31, 95.31,<br>104.92 | 32.38, 39.68,<br>47.24 | 44.02, 39.86,<br>46.26 | 38.38, 38.38,<br>59.43 |
| Unit cell Angles (α,β,γ) ° | 90, 90, 90 | 90, 90, 90 | 90, 90, 90 | 90, 98.73, 90 | 90, 90, 90 |
| I/σ (last shell) | 21.2 (0.9) | 18.7 (0.8) | 12.5 (2.0) | 26.2 (1.1) | 21.1 (2.0) |
| <sup>1</sup>R<sub>sym</sub> (last shell) | 0.082 (2.61) | 0.058 (2.884) | 0.068 (0.424) | 0.059 (0.637) | 0.039 (0.732) |
| <sup>2</sup>R<sub>meas</sub> (last shell) | 0.085 (2.723) | 0.06 (3.012) | 0.074 (0.516) | 0.070 (0.784) | 0.043 (0.809) |
| <sup>3</sup>CC<sub>1/2</sub> (last shell) | 1 (0.529) | 0.999 (0.443) | 0.997 (0.814) | 1 (0.631) | 0.999 (0.830) |
| Completeness (last shell) % | 100.0 (99.8) | 99.8 (98.1) | 80.8 (11.8) | 99.5 (98.8) | 99.8 (98.3) |
| Number of reflections | 839884 | 1388162 | 144010 | 50378 | 74958 |
| *unique* | 63659 | 106052 | 25495 | 15936 | 12610 |
| Multiplicity (last shell) | 13.2 (12.4) | 13.1 (11.7) | 5.6 (3.0) | 3.2 (2.8) | 5.9 (5.5) |
| **Refinement** |  |  |  |  |  |
| Resolution (Å) | - | 40-1.35 | 30.4-1.02 | 45.7-1.75 | 32.2-1.26 |
| No. of reflections | - | 105839 | 25448 | 15917 | 12560 |
| *working* | - | 102934 | 24251 | 15208 | 11966 |
| *free* | - | 2905 | 1197 | 709 | 594 |
| <sup>4</sup>R<sub>work</sub> (last shell) (%) | - | 16.21 (42.68) | 16.30 (17.77) | 20.98 (28.61) | 19.13 (22.50) |
| <sup>4</sup>R<sub>free</sub> (last shell) (%) | - | 16.79 (43.92) | 17.64 (17.00) | 23.70 (31.23) | 21.36 (24.19) |
| **Structure/Stereochemistry** |  |  |  |  |  |
| No. of atoms | - | 4741 | 1166 | 2531 | 642 |
| *solvent* | - | 343 | 87 | 60 | 46 |
| *ligand* | - | 0 | 10 | 0 | 0 |
| *hydrogen* | - | 2159 | 536 | 1212 | 297 |
| r.m.s.d. bond lengths (Å) | - | 0.008 | 0.008 | 0.010 | 0.011 |
| r.m.s.d. bond angles (°) | - | 0.94 | 0.993 | 1.000 | 0.896 |
| Ramachandran<br>favored/allowed (%) | - | 98.2%/100% | 100%/100% | 99.3%/100% | 100%/100% |
| Molprobity score | - | 0.82 | 0.78 | 0.64 | 1.31 |
| <sup>5</sup>SBGrid Data Bank ID | 865 | 864 | 866 | 867 | 868 |
| <sup>6</sup>Protein Data Bank ID | - | 7T5T | 7T5U | 7T5W | 7T5V |

<sup>1</sup>R<sub>sym</sub> = $\sum_j |I_j - \langle I \rangle| / \sum_j I_j$ , where $I_j$ is the intensity measurement for reflection $j$ and $\langle I \rangle$ is the mean intensity for multiply recorded reflections.
where $I_{hj}$ is a single intensity measurement for reflection $h$ , $\langle I_h \rangle$ is the average intensity measurement for multiply recorded reflections, and $n$ is the number of observations of reflection $h$ .
<sup>3</sup>CC<sub>1/2</sub> is the Pearson correlation coefficient between the average measured intensities of two randomly-assigned half-sets of the measurements of each unique reflection. CC<sub>1/2</sub> is considered significant above a value of ~0.15.
<sup>4</sup>R<sub>work, free</sub> = $\sum | |F_{\text{obs}}| - |F_{\text{calc}}| | / |F_{\text{obs}}|$ , where the working and free $R$ -factors are calculated using the working and free reflection sets, respectively.
<sup>5</sup>Diffraction data for each structure have been deposited with the SBGrid Data Bank (https://data.sbgrid.org) with the noted accession codes.
<sup>6</sup>Coordinates and structure factors for each structure have been deposited with the Protein Data Bank (http://www.rcsb.org) with the noted accession codes.

